# Land-use and forest floor explain prokaryotic metacommunity structuring and spatial turnover in Amazonian forest-to-pasture conversion areas

**DOI:** 10.1101/2020.12.30.424814

**Authors:** Fernando Igne Rocha, Thiago Gonçalves Ribeiro, Marcelo Antoniol Fontes, Stefan Schwab, Marcia Reed Rodrigues Coelho, José Francisco Lumbreras, Paulo Emílio Ferreira da Motta, Wenceslau Geraldes Teixeira, James Cole, Ana Carolina Borsanelli, Iveraldo dos Santos Dutra, Adina Howe, Aline Pacobahyba de Oliveira, Ederson da Conceição Jesus

## Abstract

Advancing extensive cattle production shifts the forest landscape and is considered one of the main drivers against biodiversity conservation in the Brazilian Amazonia. Considering soil as an ecosystem it becomes vital to identify the effects of land-use changes on soil microbial communities, structure, as well as its ecological functions and services. Herein, we explored relationships between land-use, soil types and forest floor (i.e., association between litter, root layer and bulk soil) on the prokaryotic metacommunity structuring in the Western Amazonia. Sites under high anthropogenic pressure were evaluated along a gradient of ± 800 km. Prokaryotic metacommunity are synergistically affected by soil types and land-use systems. Especially, the gradient of soil fertility and land-use shapes the structuring of the metacommunity and determines its composition. Forest-to-pasture conversion increases alpha, beta, and gamma diversities when considering only the prokaryotes from the bulk soil. Beta diversity was significantly higher in all forests when the litter and root layer were taken into account with the bulk soil. Our argumentation is that the forest floor harbors a prokaryotic metacommunity that adds at the regional scale of diversity a spatial turnover hitherto underestimated. Our findings highlight the risks of biodiversity loss and, consequently, the soil microbial diversity maintenance in tropical forests.

## 1. INTRODUCTION

Habitat fragmentation and land-use intensification have led to an alarming and rapid decline of biodiversity in tropical rainforests (Nobre et al., 2016). Soil microbiomes, which are key to ecosystem functioning and comprise a great capacity to reflect the impact of the land-use intensification on natural resources (Barnes et al., 2017), are one of the affected components of this biodiversity (Aponte et al., 2013; Ushio et al., 2010). Consequently, it is crucial to understand how the conversion of tropical forest to other land-use systems affects edaphic microbiota, especially prokaryotes (Hug et al., 2016). Previous studies have identified a strong relationship between microbial biodiversity, soil properties, and land management in the Amazon rainforest (de Carvalho et al., 2016; Jesus et al., 2009; Mendes et al., 2015; Navarrete et al., 2015; Pedrinho et al., 2019; Rodrigues et al., 2013). These findings have shown that the deforestation followed by the introduction of pastures and agricultural systems increase the alpha diversity of soil bacteria, contrary to the previous expectation that bacterial diversity would be positively correlated with plant diversity. Moreover, these studies have mostly shown that pH is one of the main abiotic factors shifting community structure in this process, hence: richness, diversity, and dominance in “local” alpha diversity.

A still unresolved question is whether intensification of converted tropical ecosystems may contribute to soil microbial homogenization across space (Petersen et al., 2019). Available studies suggests that, although land-use intensification tends to increase microbial alpha diversity, this effect does not persist on the beta diversity scale, possibly decreasing the gamma “regional” diversity and a decline in microbial turnover across space (Goss-Souza et al., 2017; Mendes et al., 2015; Rodrigues et al., 2013). However, contrasting results (de Carvalho et al., 2016; Lee-Cruz et al., 2013) indicate higher components of diversity (alpha, beta, and gamma) over more intensive land uses due to the increased environmental heterogeneity, evidencing trends contrary to microbial homogenization after the land-use change.

Previous studies have been carried out with a low variety of soil types, which reduces the ability to predict different drivers in the structuring of microbial communities, besides being predominantly limited to the surface layer of the pedosphere (i.e., borderline range between soil profile and its top organic layers). Nonetheless, organic horizons are known to sustain ecosystem functioning, especially in tropical forests (Sayer and Tanner, 2010) that predominantly grow on nutrient-poor soils (Grubb, 1995). Some recent efforts have investigated how microbial communities in the litter interact with the soil microbiota (Buscardo et al., 2018; Ritter et al., 2018), but it is still unknown how microbial communities in the forest floor (association between litter, root layer, and bulk soil) respond to regional scales of diversity (beta and gamma). Filling this gap of knowledge is important to measure the impacts of biodiversity loss in tropical forests.

In this study, we tackled how prokaryotic metacommunity (i.e., microbiomes assemblages from spatially different sites) in forest floors of the Western Amazonian forest contribute to spatial turnover and gamma “regional” diversity. In addition, forests and pastures under different soil types were also analyzed to detect variations based on the chemical and physical attributes of those soils. Our hypothesis is that the soil type rather than the land-use history is the main factor structuring prokaryotic metacommunity. Especially, we hypothesize that the inclusion of organic components of the forest floor increases overall prokaryotic metacommunity heterogeneity at alpha, beta, and gamma diversity scales. To investigate these effects, we targeted 16S rRNA through gene amplicon sequencing to assess microbiomes in a geographic gradient that covers an extensive range of soils and landscapes in the Brazilian Western Amazonian under the effects of recent forest-to-pasture conversion.

## 2. MATERIALS AND METHODS

### 2.1 Sampling and experimental design

This study was carried out in the Brazilian Western Amazonia, within a geographical range of ± 800 km, which covers spots near the cities of Bujari (state of Acre), Boca do Acre and Manicoré (state of Amazonas) (Supplementary Fig. S1). The sites were selected based on their importance for tropical forest conservation as well as the rapid advance of livestock production, which has been reported as one of the main drivers of deforestation. Sampling took place in August 2017 following the experimental design used by the Sustainable Amazonia Network (Gardner et al., 2013), with a total of 65 sampling points distributed among five forests and eight pasture areas (Supplementary Table S1). We used linear transects that were 200 m in length, including five sample points separated by 50 m from each other.

Traditional sampling for molecular microbial ecology studies usually removes the litter before sampling (de Carvalho et al., 2016; Khan et al., 2019; Mendes et al., 2015; Pedrinho et al., 2019). Nevertheless, when visiting our study sites, we observed that the forest floor has a root layer on top of the mineral soil core and right below the litter. This layer was thicker and closer to an H horizon (Fig. 1) in some forests, such as in Santo Antônio do Matupi. For this reason, we stratified samples in the forest among the litter (leaves, mostly), root layer, and mineral bulk soil (soil A-horizon at a depth of 0−10 cm; hereafter bulk soil). Only the bulk soil (0−10 cm) was sampled in pastures since no root layer nor a significant litter component existed in these systems. The forest root layer was involved by fragmented organic matter, which was the portion recovered by sieving (2 mm mesh) and used for DNA extraction. All material sampled for molecular analysis was immediately packed in sterile pouches and refrigerated at −80°C in the shortest time possible.

**Figure 1.**
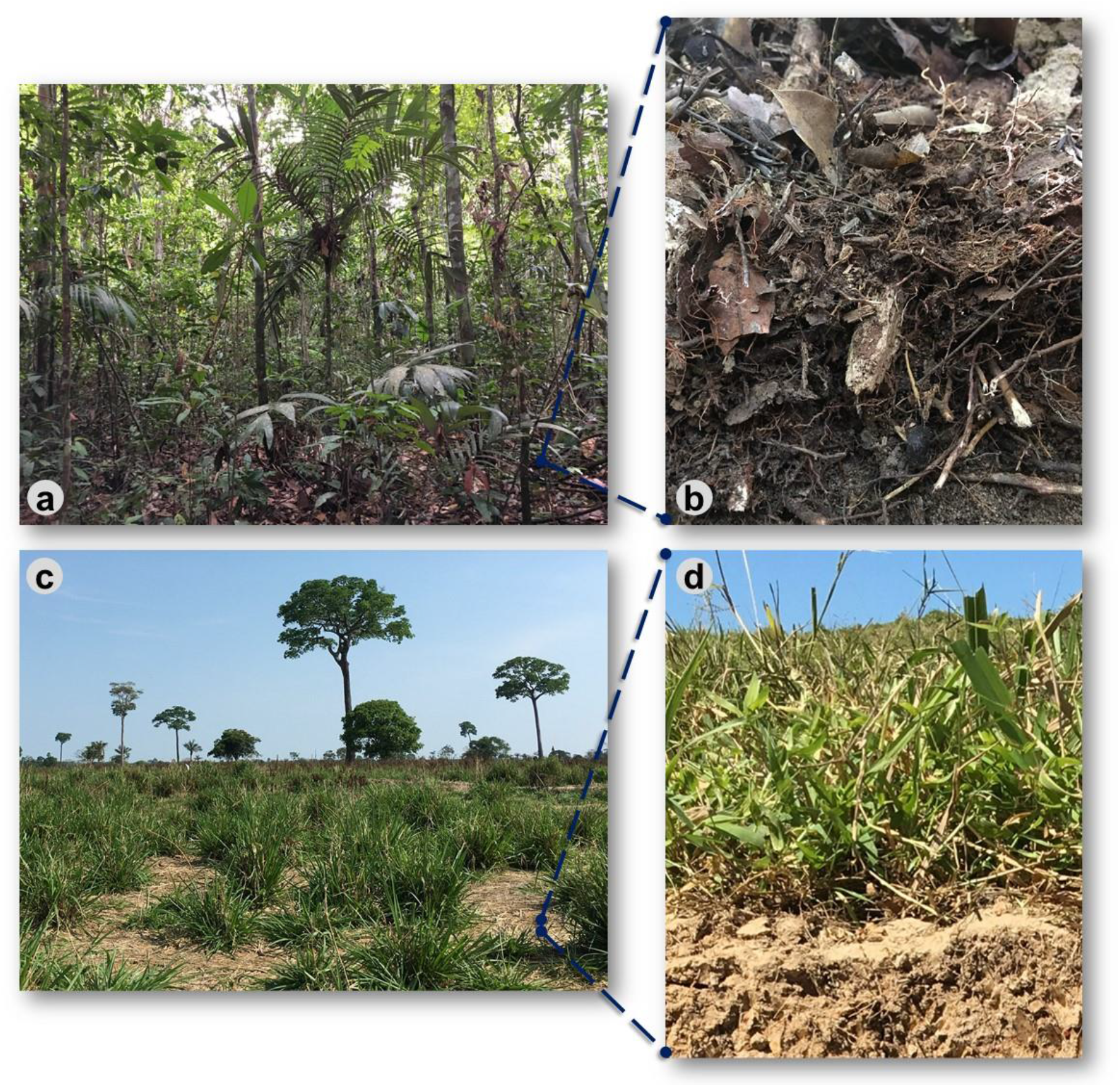
Illustrative representation of the land uses evaluated in the Brazilian Western Amazon, focusing on the decay of the forest floor after converting the forest to pasture. **a)** Rainforest where the **b)** forest floor (litter and the root layer on top of the mineral bulk soil) were sampled; **c)** Pasture systems and **d)** their respective soil surface with reduced presence of organic layers.

### 2.2 Chemical and physical analysis

Soil classification was performed for all evaluated sites, using one profile per transect where pedological description and horizon soil sampling were carried out (FAO, 2014; dos Santos et al., 2018; Santos et al., 2005). Soil physical (particle size distribution and flocculation degree) and chemical soil attributes (pH, total N, C, P, Ca^+2^, Mg^+2^, Al^+3^, potential acidity (H+Al^+3^), electrical conductivity, salts (Na^+^ and K^+^), effective CEC, as well as the calculation of derived correlations, i.e., base saturation index (%BS), base sum and Al^+3^ saturation index were analyzed (Teixeira et al., 2017) at the National Soil Research Center, Brazil. The litter was properly ground and homogenized to quantify the N and C contents using CHN elemental analysis, besides the extraction of polyphenols and tannin content following the Tropical Soil Biology and Fertility protocol (Anderson and Ingram, 1993) and conducted at the National Agrobiology Research Center, Brazil.

### 2.2 DNA extraction and high-throughput sequencing

DNA extraction from the litter, root layer, and bulk soil was performed using the standard DNeasy PowerSoil kit protocol (MO BIO Laboratories Inc.), with adjustments in the time and beating intensity of the initial protocol step after adding material to the tubes containing the beads and solution C1 (FastPrep FP120-Thermo Savant BIO101; time = 40 sec; beating = 4x). Litter and the fragmented material involving root layer samples (previously sieved in a 2 mm mesh) were macerated in liquid N with pre-sterilized mortar and pestle, as well as maintained for a minute in a water bath. Amplification of the 16S rRNA gene from all compartments was performed using barcoding DNA (Caporaso et al., 2012) with specific modifications to primer degeneracy 515F as described in (Parada et al., 2016). PCR products were purified and subjected to library preparation and sequencing with Illumina MiSeq technology following the Earth Microbiome Project protocol at the Argonne National Lab Core Sequencing Facility, USA.

### 2.3 Sequencing data processing

Sequence separation was performed in a Python environment, based on primer barcodes. The 16S rRNA sequence data were further processed, aligned, and categorized using the DADA2 microbiome pipeline (https://github.com/benjjneb/dada2) by recommended parameters with quality filtering of sequence length over 250 base pairs (Callahan et al., 2016). DADA2 characterizes microbial communities by identifying the unique amplicon sequence variants (ASVs) among the 16S rRNA reads. ASVs exhibit fewer false-positive taxa and reveal cryptic diversity, otherwise undetected by traditional OTU approaches (Callahan et al., 2017). Further, the taxonomy was assigned for each ASV assessing the Silva taxonomic training (database v132) (Quast et al., 2012). R packages ‘dada2’ v.1.14.0 (Callahan et al., 2016) and ‘decipher’ v.2.14. (Wright et al., 2012) were used in the R 3.6.1 environment (R Team, 2018).

### 2.4 Prokaryotic metacommunity analysis and environmental variable selection

The quality step (filtering, denoising, and the removal of chimeras) on the abundance matrices was used to eliminate low prevalence sequences, as well as sequences from Chloroplast, Eukaryota, and Mitochondria. After that, 2,735 ASVs were removed, resulting in a total of 1,901,440 read counts, which were divided into 15,335 ASVs with 15,221 average counts per sample. Abundances were equalized by the median sequence depth (15,212 paired-reads) to reduce abundance variability. For soil variable selection, principal component analysis (PCA) was applied on the correlation matrix to obtain a smaller subset of soil variables based on their component loadings, using ‘factoextra’ v.1.0.7 R package (Kassambara and Mundt, 2018). Non-metric multidimensional scaling (NMDS) was performed to visualize similarities among communities by factors (sites, land-use, and soil variables). The ecological distance was calculated with the Bray-Curtis dissimilarity matrix. Subsequently, the factors were compared through permutational analysis of variance (PERMANOVA) using Hellinger transformed data (Legendre and Gallagher, 2001) both with 10,000 permutations. A generalized additive model with an extra penalty (γ = 1.4) was fitted to explain the importance of each selected variable on the abundance matrix, with maximum-likelihood as a smoothing parameter estimation method (Marra and Wood, 2011). Biotic and abiotic ordinations were matched using Procrustes analysis (Peres-Neto and Jackson, 2001) that allow measuring how much they are correlated. We used differential heat-tree to visualize significant differences in taxonomic composition between the forest floor compartments in a pairwise Wilcoxon rank-sum test comparison using the ‘metacoder’ v.0.3.3 R package (Foster et al., 2017). Analyses were carried out in R environment, mainly supported by ‘phyloseq’ v.1.30.0 (McMurdie and Holmes, 2013), ‘vegan’ v.2.5-6 (Oksanen et al., 2016), and ‘ampvis2’ v.2.5.5 (Andersen et al., 2018) packages and dependencies. Finally, linear discriminant analysis (LDA) effect size (LEfSe) (Segata et al., 2011) was accessed on MicrobiomeAnalyst (Chong et al., 2020) to incorporate statistical significance with biological consistency (effect size) estimation, in a non-parametric factorial Kruskal-Wallis sum-rank test to identify features with significant differential abundance. Features with at least 2.0 log-fold changes and α < 0.05 were considered significant. All p-values were corrected by the false discovery rate method (Benjamini and Hochberg, 1995) to avoid the inflation of Type-I error due to multiple tests.

### 2.5 Diversity partitioning (α, β and γ)

HCDT entropy has been proven as a powerful tool for measuring diversity by generalizing classical indices (Marcon et al., 2014). Here, it was turned into Hill numbers, which generate effective numbers of equally frequent species for each value of “q” in a unified framework, making possible the straightforward interpretation and comparison (Chao et al., 2014). The order of diversity “q” attaches different sensitivity to rare species, being: “q = 0” the most sensitive (species richness); “q = 1” all individuals are equally weighted (exponential of Shannon’s entropy); and “q = 2” is sensitive to the dominant species (inverse of Simpson index) (Jost, 2006). Because Hill numbers are continuous and have a common unit, they can be portrayed on a single graph as a function of “q”, leading to a “diversity profile” of effective species. Further details can be found in Chao et al. (2014). Diversity partitioning means that, in a given area, the γ-diversity of all individuals found can be divided internally (α-diversity) and between the local assembly (β-diversity) (Daly et al., 2018) and was calculated for all compartments of the forest floor and pasture bulk soil. Kruskal-Wallis test was used in univariate comparisons based on the overall effective numbers (i.e., Hill’s q 0, 1 and 2) as a single way to highlight the contribution of each compartment and all the forest floor at a given diversity scale. Analyses were performed using the ‘entropart’ v.1.6-1 R package (Marcon and Hérault, 2015) and ‘stats’ v.3.6.1 (R statistical functions).

## 3 RESULTS

### 3.1 Gradient of soil fertility drives soil prokaryotic metacommunity structuring

A principal component analysis on the selected soil variables (i.e., pH, BS%, aluminum saturation, Ca + Mg, base sum, and silt; Supplementary Fig. S2) revealed 83% and 11.2% of the explained variance on PC1 and PC2, respectively. For the extracted soil variables, no statistical differences were found between the forest and pasture of BUJ, as well as between the pastures of BAC and MAN (Supplementary Table S2).

The structure of prokaryotic metacommunity (i.e., microbiome assemblages from spatially different sites) differed among the study sites as well as between land uses (PERMANOVA, F = 8.20, p < 0.001) as well as between land uses (F = 11.07, p < 0.001) for all pairwise comparisons (Supplementary Table S3). Metacommunity structure was significantly correlated to the base saturation index, showing that it shifted along a gradient of soil fertility (Fig. 2; F = 9.93, p < 0.001), from places with highly weathered soils (BAC and MAN forests) to those with high natural fertility (BUJ forest and pasture). We detected a significant statistical interaction between sites and land-use (F = 3.97, p < 0.001; Supplementary Table S3) which indicates that both factors contribute to prokaryotic community structuring, influenced by the soil type by each site and land-use characteristics, as shown further.

**Figure 2.**
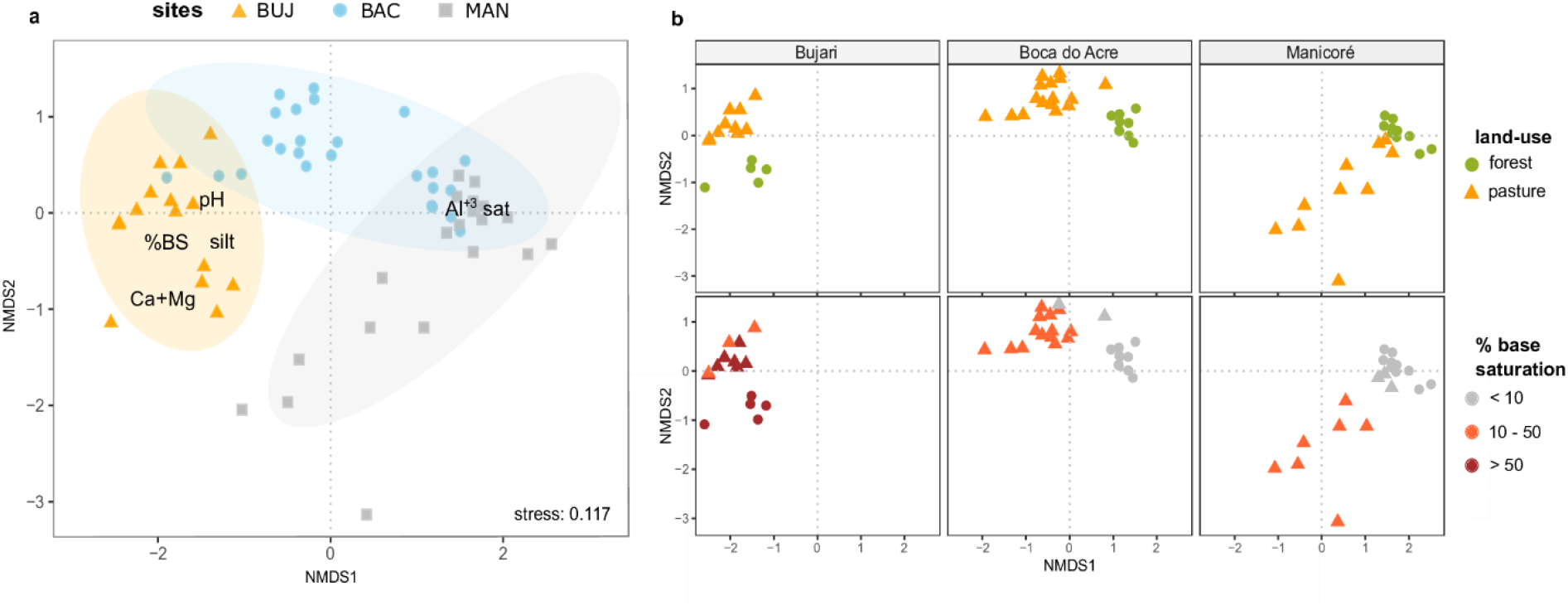
Land-use and soil type shape prokaryotic metacommunity structure in the bulk soils. **a)** Nonmetric multidimensional scaling (NMDS) based on Bray–Curtis dissimilarity among samples in the normalized ASV data of soil prokaryotic communities, highlighting the study sites and soil variables correlated with community structure; **b)** NMDS based on ecological distance (Bray-Curtis) of soil prokaryotic communities of each study area, highlighting the sample distribution pattern by land-use (upper boxes) and gradient of fertility (below boxes, by the base saturation index).

Procrustes analysis identified a positive correlation between biotic and abiotic matrices (71.83%, p < 0.001). Generalized additive models for each extracted soil variable in the PCA revealed high deviance explained for those variables, determining its importance in mediating the distribution of prokaryotic communities (Supplementary Table S4). Moreover, soil pH was positively associated with ASV richness (Supplementary Fig. S3).

### 3.2 Land-use and soil type shape the predominant composition among soil prokaryotic communities

Features that most likely explain differences between land-use systems and sites were determined by discriminant analysis (LDA) effect size (LEfSe), and patterns were detected showing taxa associated with land-use regardless of soil type. At the phylum level, *Proteobacteria, Gemmatimonadetes, Thaumarchaeota, Rokubacteria*, and *WPS-2* were revealed as the most abundant in forest systems (Supplementary Fig. S4). In contrast, *Actinobacteria, Chloroflexi, Firmicutes*, and *Bacteroidetes* were the most important phyla in pasture systems. For BAC, we found eight significantly more abundant phyla in pasture soils and four in the forest. Both BUJ and MAN had the same number of predominant phyla among their land uses. When sites were compared by the same land-use, we observed that the BUJ forest hosts the largest number of predominant phyla compared to other sites. *Verrucomicrobia*, in BUJ, and *Acidobacteria*, in BAC, are the most prevalent phyla in pasture and forest soils, respectively (Supplementary Fig. S5).

### 3.3 The structure and composition of prokaryotic metacommunity in the forest floor reflect land-use as a biotic selector

Prokaryotic metacommunity structure differed significantly among the litter, the root layer, and this result was consistent among all studied sites (Fig. 3; PERMANOVA, F = 18.08, p < 0.001). The prokaryotic metacommunity structure of the litter communities contrasted with those found in other compartments of the forest floor (Supplementary Table S5). Differences in the prokaryotic metacommunity among sites were associated with variations in litter chemical composition (Procrustes analysis: 63.2%, p < 0.001), mainly due to the polyphenol content, N content, and C:N ratio (Supplementary Fig. S6). All forest floor compartments were compared among themselves and with the pasture bulk soil. Taxa that were enriched or reduced were identified (Fig. 4, 5). *Chloroflexi, Proteobacteria, Firmicutes*, and *Verrucomicrobia* were the most statistically different (LDA; p < 0.001). *Proteobacteria* was the only phylum present in all forest compartments, especially in the litter (> 60% relative abundance; p < 0.001, LDA = 3.6). These patterns were found to be similar in all sites. *Planctomycetes* were the most representative group in the forest root layer (p < 0.001, LDA = 2.05) despite their low relative abundance (Fig. 3c). Overall, 30.2% of ASVs are shared among the compartments of the forest floor; 22.6% between BAC and MAN; 13.1% between BAC and BUJ, and only 1.3% between BUJ and MAN. BUJ has 1491 (14.2%) restrict ASVs in its microbial communities (Supplementary Fig. S7), reflecting the distinct chemical composition in the compartments of the forest floor in relation to the other sites.

**Figure 3.**
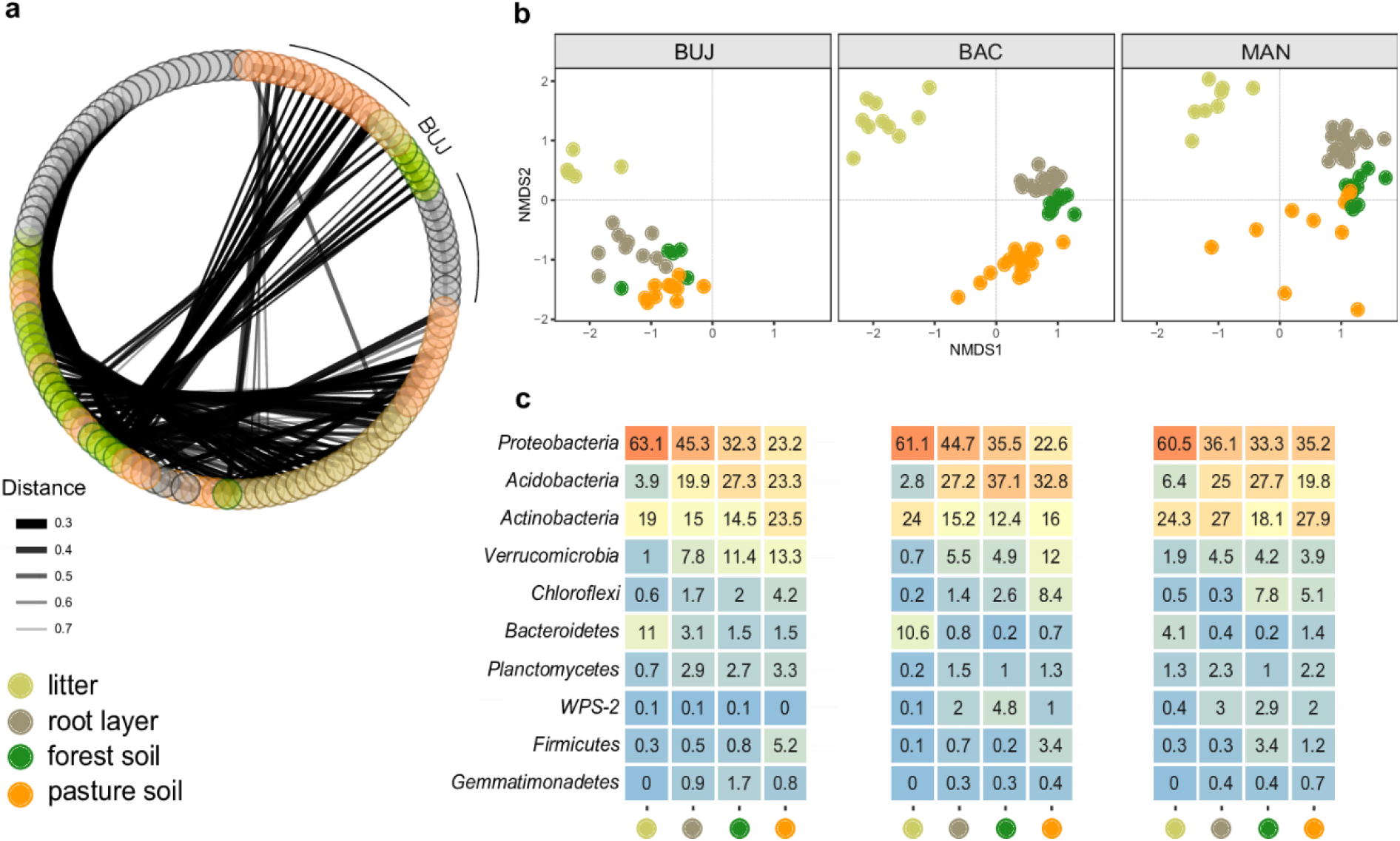
Co-occurrence of prokaryotic metacommunity between sites based on forest floor compartments and pasture bulk soil. **a)** Co-occurrence based on compartments (litter, root layer, forest and pasture soil) and study sites; the thickness of the links is proportional to the strength of the interactions; **b)** NMDS by compartments and sites (BUJ, BAC and MAN); distance measured by Bray-Curtis based on the abundance of ASVs from each sample point; **c)** Relative abundance of Bacteria (phylum level) in compartments of forest floors and pasture soil.

**Figure 4.**
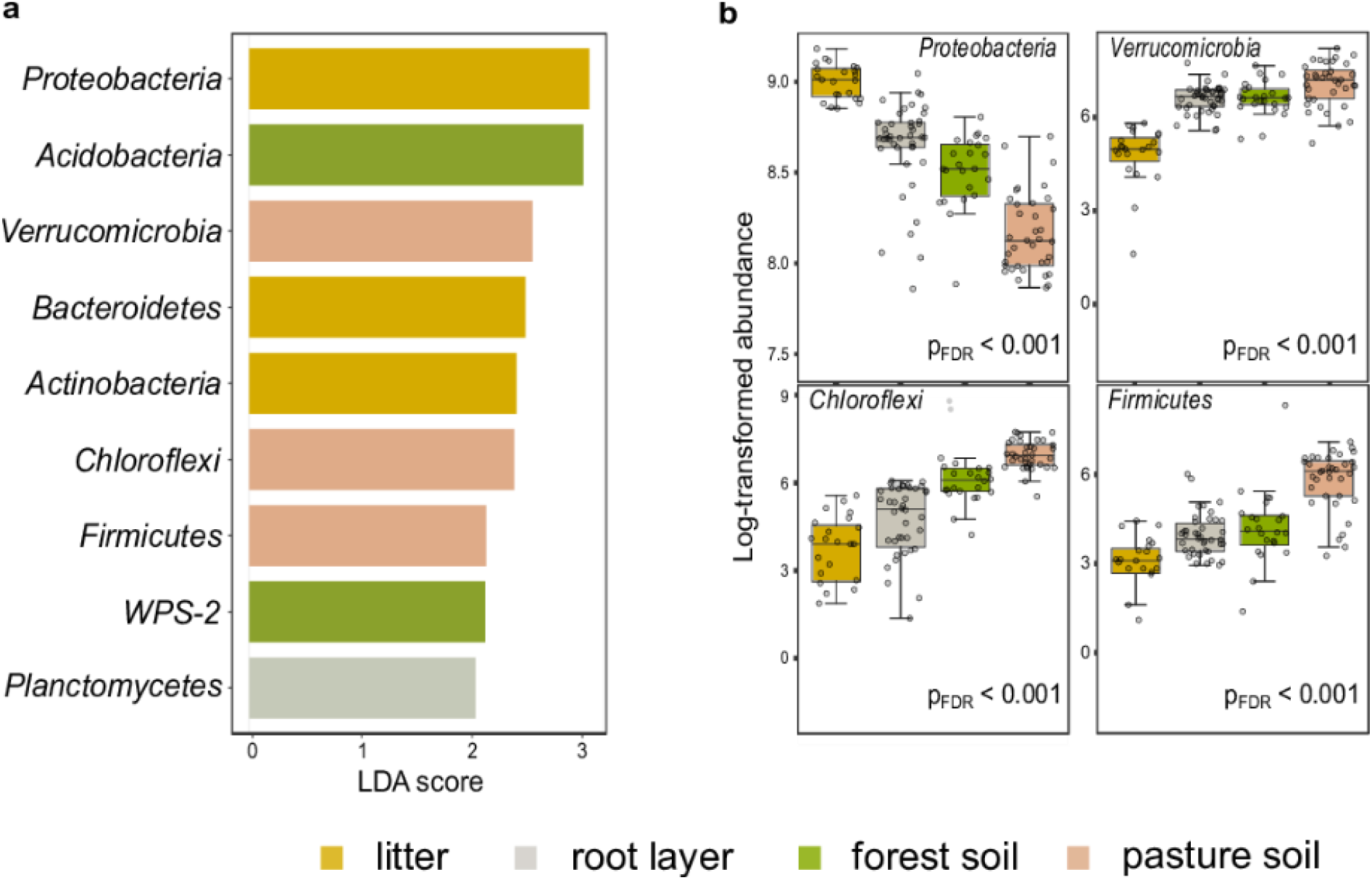
Differential abundance among the most relevant taxa in the forest floor and pasture bulk soil in the Western Brazilian Amazon. LEfSe multivariate analysis to significant differential abundances (false discovery rate adjusted p-value (pFDR) < 0.001) with LDA > 2.0; **a)** Features selected between the compartments of the forest floor and the pasture bulk soil; **b)** First four features based on pFDR < 0.001, without the application of the LDA.

**Figure 5.**
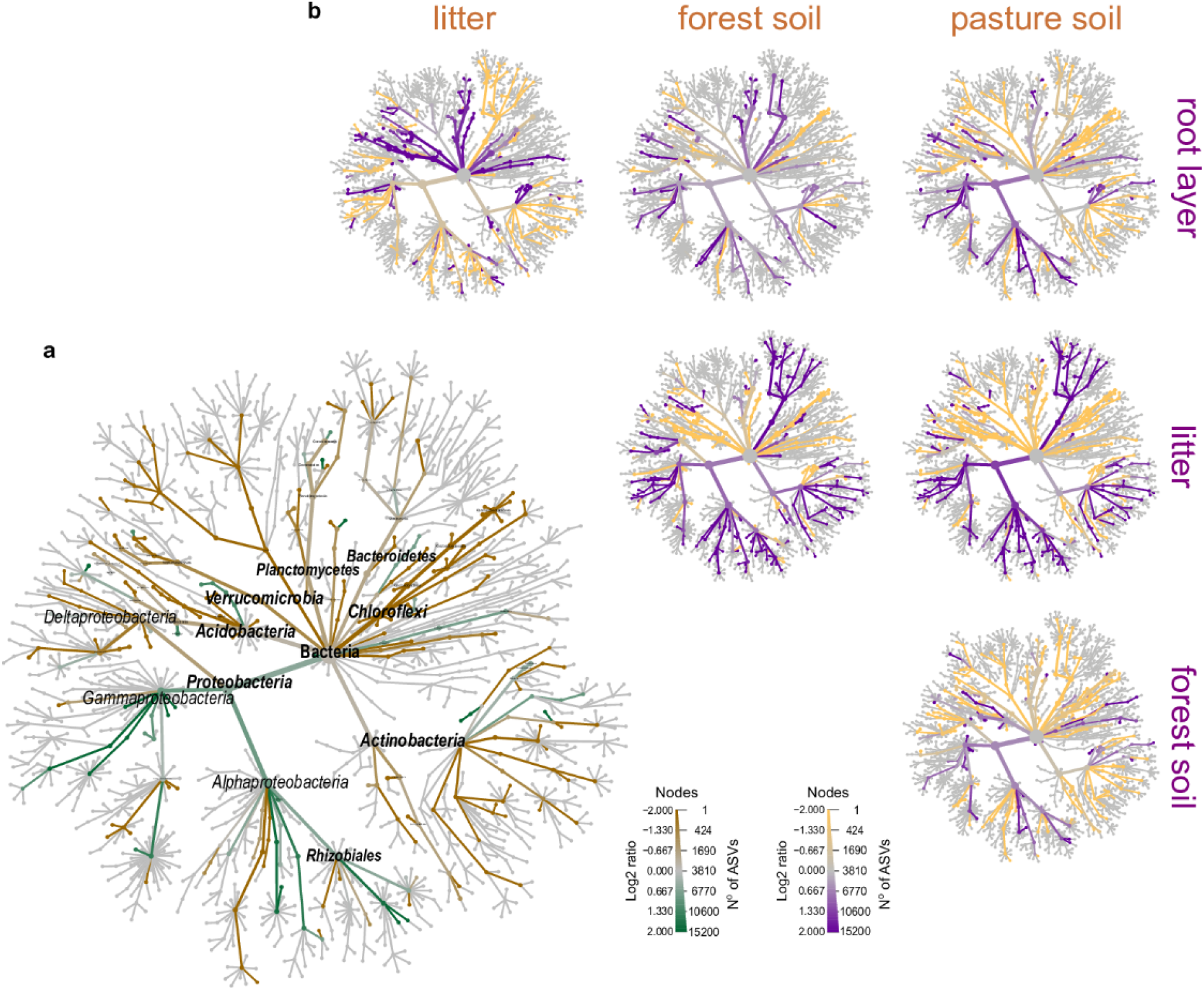
Phylogenetic differential heat tree highlighting the most expressive features among the compartments of the forest floor. **a)** Predominance of phylogenetic groups in forest floor (green color) and pasture bulk soil (brown color); **b)** Pairwise comparison between each compartment; The color of each branch represents the log-10 ratio of median proportions of reads observed at each compartment. Only significant differences are colored, determined using a Wilcox rank-sum test followed by a Benjamini-Hochberg (FDR) correction for multiple comparisons.

### 3.4 Forest floor reveals prokaryotic diversity and spatial turnover in Brazilian Western Amazonia

Diversity partitioning analysis showed that the ASV richness (Hill’s q = 0) in bulk soils is significantly higher in pastures than forests for all diversity scales and study sites, especially for MAN (Fig. 6). Beta (χ^2^ = 6.94, p < 0.001). Gamma diversity (χ^2^ = 5.43, p = 0.013) was also higher in pasture bulk soil, except for BUJ (p > 0.05). The effective number of dominant ASVs was similar (Hill’s q = 2) for any diversity scale, as well as in the comparison between forests and pastures, meaning that both systems have a similar number of dominant groups in the bulk soil. Nevertheless, when the forest floor was taken as a whole, that is, when the metacommunity in the litter, root layer, bulk soil and were taken into account, we observed that the differences in “local” alpha diversity between forest and pastures are no longer observed (Fig. 6; Supplementary Table S7). Only BAC showed a statistically higher effective number of species in its pastures for all orders of diversity “q”. BUJ had the highest alpha diversity for both litter, root layer, and bulk soils in comparison with the other study sites. Especially, the ASV richness (q = 0), as well as Shannon diversity (q = 1) and Simpson dominance (q = 2) of the forest floor showed the highest beta diversity for all study sites, reflecting increases in ASV spatial turnover. For the “regional” gamma diversity, only the forest floor of BUJ had a significant difference in the effective number of species between forest and pasture (χ^2^ = 6.64, p = 0.009).

**Figure 6.**
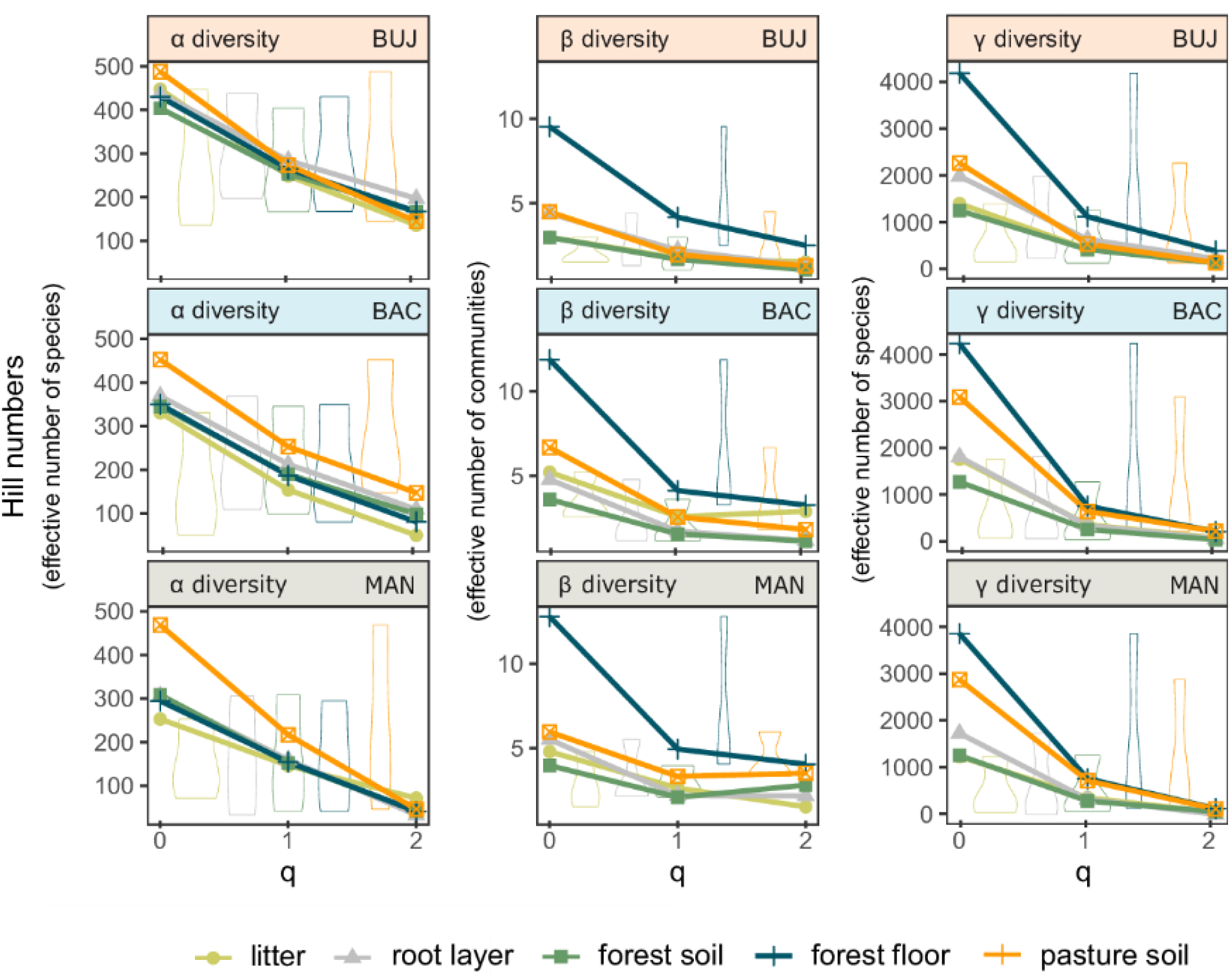
Diversity partitioning analysis evidencing the heterogeneity of prokaryotic metacommunity across land uses. Alpha (α), beta (β) and gamma (γ) (i.e., local, community, and regional) diversities for the forest and pasture bulk soils, litter, root layer, and the forest floor in each site. Hill numbers (q = 0, ASV richness), (q = 1, exponential of Shannon’s entropy for equally weighted ASVs) and (q = 2, inverse of Simpson index for dominant taxa).

## 4 DISCUSSION

In the past years, several authors have addressed the issue of increasing microbial alpha diversity among anthropized systems, such as mechanized agriculture and pastures (de Carvalho et al., 2016; Lammel et al., 2018; Rodrigues et al., 2013). This topic has been highlighted because more diversified communities in more intensified areas contradict the common sense that high diversity should necessarily be related to pristine environments. Here, we proposed an integrated view of the forest floor in which the litter and root layer were also considered as an integral part of the forest soil ecosystem (Ponge, 2015). These two components are a source of nutrients to soil microorganisms, where plant roots and microbiota are more active, rather than in bulk soil (Biswas and Kole, 2017; Gobat et al., 2004). Nevertheless, except for a few studies (Buscardo et al., 2018; Ritter et al., 2018), the forest floor was not considered as a whole unit in the diversity surveys carried out in the Amazonia (de Carvalho et al., 2016; Jesus et al., 2009; Mendes et al., 2015; Navarrete et al., 2015; Pedrinho et al., 2019; Rodrigues et al., 2013). For this reason, our perception was that the microbial diversity of the forest soil has been hitherto underestimated. This observation gains more relevance when we consider that the litter and the root layer are the soil compartments most affected by deforestation. Clearing techniques traditionally used to remove the forest involve the burning of the most part of its biomass, which is the major method of deforestation practiced in Amazonia (Paula et al., 2014).

Thus, we sampled the litter and roots present in the forest floor to understand how prokaryotic communities are composed and structured in these compartments, as well as if these components of the forest floor add to the partitioning of the diversity in the forest and, consequently, how they affect the comparison with the microbial diversity found in pasture soils.

### 4.1 Prokaryotic metacommunity reflects the synergistic interaction between land-use and soil type

Multiple analyzes based on the next-generation sequencing approach have allowed us to support our first hypothesis that soil type, rather than land use patterns, mainly leads to the structuring of the prokaryotic metacommunity in bulk soils. This finding highlights that soil variables, especially those related to the soil fertility, such as pH and base saturation are the major attributes driving the prokaryotic community structuring in bulk soils.

Our argument is based on the observation that communities belonging to the most distant geographic areas (± 650 km; BAC to MAN) showed greater similarity in their structure (Bray-Curtis distance = 0.51) than communities from nearby sites (± 150 km; BUJ to BAC; Bray-Curtis = 0.87). This distinction reflects the influence of different soil-forming processes on microbial community structuring. Soils of the State of Acre mostly come from weathering sedimentary rocks, and specifically those found in this study are a patch of naturally eutrophic soil, such as Luvisols (Bernini et al., 2013). BAC and MAN have predominantly Acrisols and Ferralsols, highly weathered soils developed on sandstones and claystones, and mainly formed on remnants of ferrallitic plateaus and convex hills which are not flooded (Souza et al., 2018). The gradient of soil fertility across soils with distinct pedogenesis and weathering degrees is a major contribution of this study to understanding how microbiomes are modeled by the forest floor under the same land-use system. Soil pH may not directly alter prokaryotic community structure but may be considered as an integrating variable that provides an index of soil conditions instead (Lauber et al., 2009). Many soil attributes, such as nutrient availability, cationic metal solubility, organic C characteristics, soil moisture regimen, and salinity, are often directly or indirectly related to soil pH (Bissett et al., 2011; Suleiman et al., 2013). However, recent studies indicate that bacterial community assembly processes differ in relation to soil pH, with near-neutral pH leading to more stochastic communities, whereas extreme conditions lead to more deterministic assembly and clustered communities (Tripathi et al., 2018). Thus, the influence of variables such as temperature are mainly revealed where soil pH is relatively constant (Nottingham et al., 2018).

Our results consistently support previous studies that also showed the cause-effect relationship of the changes in soil pH and alterations in the natural structure and composition of the soil microbiomes, due to the land-use conversion (Berkelmann et al., 2018; Goss-Souza et al., 2017; Jesus et al., 2009; Mendes et al., 2015; Navarrete et al., 2015). Moreover, regarding the taxonomic approach of communities, we observe a fingerprint through land uses for some taxa, even considering the different soil types. *Actinobacteria* was dominant in the pasture system. In opposition to this, *Proteobacteria* were considerably higher in the forest floor, especially in the litter. *Proteobacteria* are usually related to high levels of organic C and have been extensively reported as an indicator of land-use change as its high abundance is drastically reduced after the conversion of the rainforest into simplified systems such as pastures or mechanized agriculture (de Carvalho et al., 2016; Mendes et al., 2015; Navarrete et al., 2015). *Proteobacteria*, specifically *Alphaproteobacteria*, and *Gammaproteobacteria*, are functionally important in natural systems that are known to undergo weak soil perturbation and provide copiotroph habitats rich in recalcitrant organic matter (Pascault et al., 2013). They are also closely related to methane oxidation (CH_4_) due to its methanotrophic characteristics, helping to mitigate the emissions of this gas by the balance between production and consumption within systems with lower anthropic disturbance, such as forests (Tate, 2015). In more disturbed systems, the soil microbiota responds by changing dominant groups, as feedback for the land-use management, which can print significant shifts in soil attributes over time. Increases in the relative abundance of *Actinobacteria* and *Chloroflexi* populations were highlighted in Fierer et al. (2012) and Mendes et al. (2015). *Actinobacteria* are functionally related to organic substrate decomposers and present the ability to produce spores, allowing this group to maintain its activity in more anthropized systems (Ventura et al., 2007). Some groups of the *Chloroflexi* are thermophilic aerobes, having the ability to develop their metabolism at high temperatures, also keeping an important relationship in the decomposition of organic matter (Yamada et al., 2005) and consequently, predominance in pasture soils.

### 4.2 Role of the prokaryotic metacommunity in the forest floor and deforestation as a risk for its maintenance

The tropical forest floor undoubtedly plays an important role in the biodiversity and ecosystem functioning on a global scale (Poorter et al., 2015). The biogeochemical cycles in that ecosystem regulate the largest terrestrial C storage, maintaining high biomass and productivity, although mostly growing on nutrient-poor soils (Finzi et al., 2011; Sayer et al., 2020). However, the rapid advance of land-use change through the livestock expansion represents a high risk for its maintenance, because the forest floor is mineralized as a short-term effect after the forest-to-pasture conversion, with no subsequent replacement of the compartments that compose it. Some efforts to evidence nutrient retention and uptake in the forest floor have been done (Sayer et al., 2020; Sayer and Tanner, 2010), considering that measurements in the mineral soil only represent a small part of the picture. Hence, a better understanding of the role of the prokaryotic communities in the forest floor, as well as how are impacted by land-use alteration is essential to predict consequences in the face of global changes (Lladó et al., 2017; Rillig et al., 2019).

Firstly, our investigation of forests in the Western Amazonia suggests that litter prokaryotes apparently do not have a direct relationship with the root layer and soil microbiota, therefore they are not directly influenced by soil attributes. It is noteworthy that litter microbiota is likely predominantly endophytic and related to the floristic composition of the forests. A specific litter quality chemically related to the forest phytophysiognomy is added to the forest floor, providing different drivers for the microbial community structuring (Buscardo et al., 2018). Nonetheless, plant diversity and community composition are influenced by geology and physicochemical soil properties (Higgins et al., 2011; Ritter et al., 2018), which is indirectly important to explain variations in composition and structure of the litter microbiome. *Proteobacteria, Actinobacteria*, and *Bacteroidetes* were the most abundant phyla for that compartment, as already evidenced by Purahong et al. (2016) and Tlàskal et al. (2016). Moreover, we observed differences in the communities between study sites, which are explained by the chemical composition of the litter. We detected a higher content of polyphenols, tannins, and C:N ratio, mostly in the litter of the MAN’s forests, which may be related to the highest relative abundance of *Actinobacteria* (Lewin et al., 2016) and a smaller abundance of *Bacteroidetes* (Xue et al., 2016) compared to the other study sites.

The root layer, especially fine-root production and turnover is a highly important input of plant detritus to the soil. It is also a key energy source to soil microbiomes, and consequently, a major pathway of nutrient flux in terrestrial ecosystems (Yuan and Chen, 2010; Zechmeister-Boltenstern et al., 2015). In our study, the root layer-associated prokaryotes also showed significant differences in the community structure between study sites but sharing similarities with the bulk soil due to its transient position on the forest floor. Despite the differences in community structure, we observed a clear pattern related to the microbiota composition, with a significant enrichment of *Planctomycetes* in the root layer for all study sites. Some planctomycetes may be involved in degrading polymeric organic matter (Ivanova et al., 2018), however, experimental data remain scarce due to the low number of characterized representatives of this phylum. The higher relative abundance of *Planctomycetes* in the root layer have already been reported for the Amazonian rainforest (Fonseca et al., 2018), nevertheless more studies are needed to understand the ecological role of planctomycetes along the root layer in tropical rainforests, as well as its potential representativity for that ecological niche.

### 4.2 Forest floor as an ecosystem for accessing microbial diversity in tropical forests

Although our results agreed with previous studies that have identified higher alpha diversity in pasture soils compared to forests, a better understanding of microbial turnover and “regional” gamma diversity is still on demand, as pointed out by Petersen et al. (2019) in a recent meta-analysis that tackled the soil microbiota in tropical land uses.

Our diversity partitioning analysis does not indicate a positive correlation between plant and soil prokaryotic beta diversity, as found by Prober et al. (2015) neither does it indicate the reduction of spatial heterogeneity in pastures introduced after deforestation, as evidenced by Rodrigues et al. (2013) in the Western Amazonia and Goss-Souza et al. (2017) in the Atlantic Rainforest. Our results agreed with the findings described by de Carvalho et al. (2016), who found a higher beta diversity for soil prokaryotes in more altered land uses of the Eastern Amazonia, such as pastures, especially for ASV richness (q = 0) and Shannon diversity (q = 1).

Nevertheless, when forest litter and root layer are taken into account with the bulk soil, we detect a higher effective number of communities (beta diversity) in all forests, rather than in the pastures. This allows us to realize that, at least in tropical forests, regardless of the soil type, the biodiversity intricate in the forest floor might confer similar abilities for the ecological functioning of the forest ecosystem, such as nutrient cycling and C sequestration. Aspects related to the functional redundancy in the forest floor is a key topic to be further investigated. Thus, when observed individually, the litter, root layer, and bulk soil compartments do not give clear information on the turnover of prokaryotic communities, so it is necessary to apply an integrated interpretation of this system. Regarding gamma diversity, BUJ was the only site with differences between the forest floor and pasture bulk soil. The non-overlapping of the gamma diversity in less sensitive values (q = 1 and q = 2; see Fig. 6) may indicate that the higher natural fertility found in bulk soils, in addition to the higher labile N content in the litter, should provide greater support for more stable prokaryotic diversity. Finally, our results partially corroborate our second hypothesis since we have not seen consistent increases in alpha “local” and gamma “regional” diversities after including forest floor in the diversity partition analysis.

## 5. CONCLUSIONS

Altogether, our results support previous studies that show significant shifts in the composition and diversity of bulk soil prokaryotic communities after forest-to-pasture conversion. By adding the forest litter and root layer to the bulk soil in our measurements, we demonstrate that prokaryotes vary in their community structure and composition among the forest floor compartments, with an important site-specific influence. Despite this, all pasture bulk soils have prokaryotes more correlated with increases in soil pH and base saturation, resulting in higher alpha, beta, and gamma diversities. Nevertheless, beta diversity was significatively high in all forests when the litter and root layer compartments were taken into account, highlighting increases in microbial heterogeneity across space. Our findings offer evidence to better understand the importance of the forest floor to comprehend the dynamics of microbial communities across tropical forests threatened by deforestation for pasture introduction, besides giving new perspectives on the issue of biotic homogenization. For future efforts, other components of the pasture floor should be characterized and included to generate a better picture of the presented scenario.

## 6. AUTHOR’S CONTRIBUTIONS

I.S.D., A.P.O., J.C., and E.C.J. designed the study. A.P.O., A.C.B., and E.C.J. conducted the sampling. F.I.R., T.G.R., S.S., M.R.R.C., and M.A.F. conducted the laboratory analyses. Soil descriptions and characterization were performed with support from A.P.O., J.F.L., P.E.F., W.G.T. Data analysis was conducted by F.I.R with support from A.C.H. Manuscript writing was led by F.I.R., A.C.H., and E.C.J. All co-authors contributed to the drafts and gave final approval for publication.

## 7. ACKNOWLEDGEMENTS

We acknowledge the USAID and the National Academies of Sciences, Engineering, and Medicine of the United States (NAS) for funding our research under PEER project 4−299, USAID agreement AID-OAA-A-11-00012. Any opinions, findings, conclusions, or recommendations expressed here are those of the authors alone and do not necessarily reflect the views of USAID or the NAS. We also thank CNPq, Brazil, for the research fellowships provided to Ederson da Conceição Jesus (project 475168/2012-7) and Fernando Igne Rocha (165571/2017-9). FIR was also supported by CAPES, Brazil (PDSE call n° 41/2018). Finally, we thank Sarah Owens and Stephanie M. Greenwald, both from the Argonne National Laboratory, for supporting us with the sequencing analysis.

## 10. DATA AVAILABILITY

DNA sequence data are accessible at the MG-RAST under the accession number 94905 (http://www.mg-rast.org/linkin.cgi?project=mgp94905).

## REFERENCES

Andersen, K.S., Kirkegaard, R.H., Karst, S.M., Albertsen, M., 2018. ampvis2: an R package to analyse and visualise 16S rRNA amplicon data. BioRxiv 299537.

Anderson, J.M., Ingram, J., 1993. Tropical Soil Biology and Fertility. A handbook of methods. CAB. International UK.

Aponte, C., García, L. V, Marañón, T., 2013. Tree species effects on nutrient cycling and soil biota: A feedback mechanism favouring species coexistence. Forest Ecology and Management 309, 36–46. doi:https://doi.org/10.1016/j.foreco.2013.05.035

Barnes, A.D., Allen, K., Kreft, H., Corre, M.D., Jochum, M., Veldkamp, E., Clough, Y., Daniel, R., Darras, K., Denmead, L.H., Farikhah Haneda, N., Hertel, D., Knohl, A., Kotowska, M.M., Kurniawan, S., Meijide, A., Rembold, K., Edho Prabowo, W., Schneider, D., Tscharntke, T., Brose, U., 2017. Direct and cascading impacts of tropical land-use change on multi-trophic biodiversity. Nature Ecology & Evolution 1, 1511–1519. doi:10.1038/s41559-017-0275-7

Benjamini, Y., Hochberg, Y., 1995. Controlling the false discovery rate: a practical and powerful approach to multiple testing. Journal of the Royal Statistical Society: Series B (Methodological) 57, 289–300.

Berkelmann, D., Schneider, D., Engelhaupt, M., Heinemann, M., Christel, S., Wijayanti, M., Meryandini, A., Daniel, R., 2018. How rainforest conversion to agricultural systems in Sumatra (Indonesia) affects active soil bacterial communities. Frontiers in Microbiology 9, 2381.

Bernini, T. de A., Pereira, M.G., Fontana, A., Anjos, L.H.C. dos Calderano, S.B., Wadt, P.G.S., Moraes, A.G. de L. Santos, L.L. dos, 2013. Taxonomia de solos desenvolvidos sobre depósitos sedimentares da Formação Solimões no Estado do Acre. Bragantia 72, 71–80.

Bissett, A., Richardson, A.E., Baker, G., Thrall, P.H., 2011. Long-term land use effects on soil microbial community structure and function. Applied Soil Ecology 51, 66–78. doi:https://doi.org/10.1016/j.apsoil.2011.08.010

Biswas, T., Kole, S.C., 2017. Soil organic matter and microbial role in plant productivity and soil fertility, in: Advances in Soil Microbiology: Recent Trends and Future Prospects. Springer, pp. 219–238.

Brown, D.S., Brown, J.C., Brown, C., 2016. Land occupations and deforestation in the Brazilian Amazon. Land Use Policy 54, 331–338. doi: https://doi.org/10.1016/j.landusepol.2016.02.003

Buscardo, E., Geml, J., Schmidt, S.K., Freitas, H., da Cunha, H.B., Nagy, L., 2018. Spatio-temporal dynamics of soil bacterial communities as a function of Amazon forest phenology. Scientific Reports 8, 4382. doi:10.1038/s41598-018-22380-z

Callahan, B.J., McMurdie, P.J., Holmes, S.P., 2017. Exact sequence variants should replace operational taxonomic units in marker-gene data analysis. The ISME Journal 11, 2639– 2643. doi:10.1038/ismej.2017.119

Callahan, B.J., McMurdie, P.J., Rosen, M.J., Han, A.W., Johnson, A.J.A., Holmes, S.P., 2016. DADA2: High-resolution sample inference from Illumina amplicon data. Nature Methods 13, 581–583. doi:10.1038/nmeth.3869

Caporaso, J.G., Lauber, C.L., Walters, W.A., Berg-Lyons, D., Huntley, J., Fierer, N., Owens, S.M., Betley, J., Fraser, L., Bauer, M., Gormley, N., Gilbert, J.A., Smith, G., Knight, R., 2012. Ultra-high-throughput microbial community analysis on the Illumina HiSeq and MiSeq platforms. The ISME Journal 6, 1621–1624. doi:10.1038/ismej.2012.8

Chao, A., Chiu, C.-H., Jost, L., 2014. Unifying species diversity, phylogenetic diversity, functional diversity, and related similarity and differentiation measures through Hill numbers. Annual Review of Ecology, Evolution, and Systematics 45, 297–324.

Chong, J., Liu, P., Zhou, G., Xia, J., 2020. Using MicrobiomeAnalyst for comprehensive statistical, functional, and meta-analysis of microbiome data. Nature Protocols 15, 799–821. doi:10.1038/s41596-019-0264-1

Daly, A.J., Baetens, J.M., De Baets, B., 2018. Ecological diversity: measuring the unmeasurable. Mathematics 6, 119.

de Carvalho, T.S., Jesus, E. da C., Barlow, J., Gardner, T.A., Soares, I.C., Tiedje, J.M., Moreira, F.M. de S., 2016. Land use intensification in the humid tropics increased both alpha and beta diversity of soil bacteria. Ecology 97, 2760–2771. doi:10.1002/ecy.1513

dos Santos, H.G., Jacomine, P.K.T., Dos Anjos, L.H.C., De Oliveira, V.A., Lumbreras, J.F., Coelho, M.R., De Almeida, J.A., de Araujo Filho, J.C., de Oliveira, J.B., Cunha, T.J.F., 2018. Sistema brasileiro de classificação de solos. Brasília, DF: Embrapa, 2018.

FAO - Food and Agriculture Organization of the United Nations, 2014. World reference base for soil resources 2014: International soil classification system for naming soils and creating legends for soil maps.

Fierer, N., Lauber, C.L., Ramirez, K.S., Zaneveld, J., Bradford, M.A., Knight, R., 2012. Comparative metagenomic, phylogenetic and physiological analyses of soil microbial communities across nitrogen gradients. The ISME Journal 6, 1007–1017.

Finzi, A.C., Austin, A.T., Cleland, E.E., Frey, S.D., Houlton, B.Z., Wallenstein, M.D., 2011. Responses and feedbacks of coupled biogeochemical cycles to climate change: examples from terrestrial ecosystems. Frontiers in Ecology and the Environment 9, 61–67.

Fonseca, J.P., Hoffmann, L., Cabral, B.C.A., Dias, V.H.G., Miranda, M.R., de Azevedo Martins, A.C., Boschiero, C., Bastos, W.R., Silva, R., 2018. Contrasting the microbiomes from forest rhizosphere and deeper bulk soil from an Amazon rainforest reserve. Gene 642, 389–397. doi:https://doi.org/10.1016/j.gene.2017.11.039

Foster, Z.S.L., Sharpton, T.J., Grünwald, N.J., 2017. Metacoder: An R package for visualization and manipulation of community taxonomic diversity data. PLoS Computational Biology 13, e1005404. doi:10.1371/journal.pcbi.1005404

Gardner, T.A., Ferreira, J., Barlow, J., Lees, A.C., Parry, L., Vieira, I.C.G., Berenguer, E., Abramovay, R., Aleixo, A., Andretti, C., Aragão, L.E.O.C., Araújo, I., de Ávila, W.S., Bardgett, R.D., Batistella, M., Begotti, R.A., Beldini, T., de Blas, D.E., Braga, R.F., Braga, D. de L., de Brito, J.G., de Camargo, P.B., Campos dos Santos, F., de Oliveira, V.C., Cordeiro, A.C.N., Cardoso, T.M., de Carvalho, D.R., Castelani, S.A., Chaul, J.C.M., Cerri, C.E., Costa, F. de A., da Costa, C.D.F., Coudel, E., Coutinho, A.C., Cunha, D., D’Antona, Á., Dezincourt, J., Dias-Silva, K., Durigan, M., Esquerdo, J.C.D.M., Feres, J., Ferraz, S.F. de B., Ferreira, A.E. de M., Fiorini, A.C., da Silva, L.V.F., Frazão, F.S., Garrett, R., Gomes, A. dos S., Gonçalves, K. da S., Guerrero, J.B., Hamada, N., Hughes, R.M., Igliori, D.C., Jesus, E. da C., Juen, L., Junior, M., Junior, J.M.B. de O., Junior, R.C. de O., Junior, C.S., Kaufmann, P., Korasaki, V., Leal, C.G., Leitão, R., Lima, N., Almeida, M. de F.L., Lourival, R., Louzada, J., Nally, R. Mac Marchand, S., Maués, M.M., Moreira, F.M.S., Morsello, C., Moura, N., Nessimian, J., Nunes, S., Oliveira, V.H.F., Pardini, R., Pereira, H.C., Pompeu, P.S., Ribas, C.R., Rossetti, F., Schmidt, F.A., da Silva, R., da Silva, R.C.V.M., da Silva, T.F.M.R., Silveira, J., Siqueira, J.V., de Carvalho, T.S., Solar, R.R.C., Tancredi, N.S.H., Thomson, J.R., Torres, P.C., Vaz-de-Mello, F.Z., Veiga, R.C.S., Venturieri, A., Viana, C., Weinhold, D., Zanetti, R., Zuanon, J., 2013. A social and ecological assessment of tropical land uses at multiple scales: the Sustainable Amazon Network. Philosophical Transactions of the Royal Society B: Biological Sciences 368, 20120166. doi:10.1098/rstb.2012.0166

Fierer, N., Lauber, C.L., Gobat, J.-M., Aragno, M., Matthey, W., 2004. The living soil: fundamentals of soil science and soil biology. Science Publishers.

Goss-Souza, D., Mendes, L.W., Borges, C.D., Baretta, D., Tsai, S.M., Rodrigues, J.L.M., 2017. Soil microbial community dynamics and assembly under long-term land use change. FEMS Microbiology Ecology 93.

Grubb, P.J., 1995. Mineral Nutrition and Soil Fertility in Tropical Rain Forests BT - Tropical Forests: Management and Ecology, in: Lugo, A.E., Lowe, C. (Eds.),. Springer New York, New York, NY, pp. 308–330. doi:10.1007/978-1-4612-2498-3_12

Lee-Cruz, L., Edwards, D.P., Tripathi, B.M., Adams, J.M., 2013. Impact of logging and forest conversion to oil palm plantations on soil bacterial communities in Borneo. Applied and Environmental Microbiology 79, 7290–7297.

Higgins, M.A., Ruokolainen, K., Tuomisto, H., Llerena, N., Cardenas, G., Phillips, O.L., Vásquez, R., Räsänen, M., 2011. Geological control of floristic composition in Amazonian forests. Journal of Biogeography 38, 2136–2149.

Hug, L.A., Baker, B.J., Anantharaman, K., Brown, C.T., Probst, A.J., Castelle, C.J., Butterfield, C.N., Hernsdorf, A.W., Amano, Y., Ise, K., Suzuki, Y., Dudek, N., Relman, D.A., Finstad, K.M., Amundson, R., Thomas, B.C., Banfield, J.F., 2016. A new view of the tree of life. Nature Microbiology 1, 16048. doi:10.1038/nmicrobiol.2016.48

Ivanova, A.A., Wegner, C.-E., Kim, Y., Liesack, W., Dedysh, S.N., 2018. Metatranscriptomics reveals the hydrolytic potential of peat-inhabiting Planctomycetes. Antonie van Leeuwenhoek 111, 801–809. doi:10.1007/s10482-017-0973-9

Jesus, E. da C., Marsh, T.L., Tiedje, J.M., Moreira, F.M. de S., 2009. Changes in land use alter the structure of bacterial communities in Western Amazon soils. The ISME Journal 3, 1004– 1011. doi:10.1038/ismej.2009.47

Jost, L., 2006. Entropy and diversity. Oikos 113, 363–375.

Kassambara, A., Mundt, F., 2018. Factoextra: Extract and visualize the results of multivariate data analyses. 2017. R Package Version 1.

Khan, M.A.W., Bohannan, B.J.M., Nüsslein, K., Tiedje, J.M., Tringe, S.G., Parlade, E., Barberán, A., Rodrigues, J.L.M., 2019. Deforestation impacts network co-occurrence patterns of microbial communities in Amazon soils. FEMS Microbiology Ecology 95, fiy230.

Lammel, D.R., Barth, G., Ovaskainen, O., Cruz, L.M., Zanatta, J.A., Ryo, M., de Souza, E.M., Pedrosa, F.O., 2018. Direct and indirect effects of a pH gradient bring insights into the mechanisms driving prokaryotic community structures. Microbiome 6, 106.

Lauber, C.L., Hamady, M., Knight, R., Fierer, N., 2009. Pyrosequencing-based assessment of soil pH as a predictor of soil bacterial community structure at the continental scale. Applied and Environmental Microbiology 75, 5111–5120.

Legendre, P., Gallagher, E.D., 2001. Ecologically meaningful transformations for ordination of species data. Oecologia 129, 271–280. doi:10.1007/s004420100716

Lewin, G.R., Carlos, C., Chevrette, M.G., Horn, H.A., McDonald, B.R., Stankey, R.J., Fox, B.G., Currie, C.R., 2016. Evolution and ecology of Actinobacteria and their bioenergy applications. Annual Review of Microbiology 70, 235–254.

Lladó, S., López-Mondéjar, R., Baldrian, P., 2017. Forest soil bacteria: diversity, involvement in ecosystem processes, and response to global change. Microbiology and Molecular Biology Reviews 81.

Marcon, E., Hérault, B., 2015. entropart: An R package to measure and partition diversity. Journal of Statistical Software 67, 1–26.

Marcon, E., Zhang, Z., Hérault, B., 2014. The decomposition of similarity-based diversity and its bias correction. HAL, hal-00989454(version 1), 1–12.

Marra, G., Wood, S.N., 2011. Practical variable selection for generalized additive models. Computational Statistics & Data Analysis 55, 2372–2387. doi:https://doi.org/10.1016/j.csda.2011.02.004

McMurdie, P.J., Holmes, S., 2013. phyloseq: an R package for reproducible interactive analysis and graphics of microbiome census data. PloS One 8, e61217. doi:10.1371/journal.pone.0061217

Mendes, L.W., de Lima Brossi, M.J., Kuramae, E.E., Tsai, S.M., 2015. Land-use system shapes soil bacterial communities in Southeastern Amazon region. Applied Soil Ecology 95, 151– 160.

Mendes, L.W., Tsai, S.M., Navarrete, A.A., de Hollander, M., van Veen, J.A., Kuramae, E.E., 2015. Soil-borne microbiome: linking diversity to function. Microbial Ecology 70, 255–265. doi:10.1007/s00248-014-0559-2

Navarrete, A.A., Tsai, S.M., Mendes, L.W., Faust, K., de Hollander, M., Cassman, N.A., Raes, J., van Veen, J.A., Kuramae, E.E., 2015. Soil microbiome responses to the short-term effects of Amazonian deforestation. Molecular Ecology 24, 2433–2448. doi:10.1111/mec.13172

Nobre, C.A., Sampaio, G., Borma, L.S., Castilla-Rubio, J.C., Silva, J.S., Cardoso, M., 2016. Land-use and climate change risks in the Amazon and the need of a novel sustainable development paradigm. Proceedings of the National Academy of Sciences of the United States of America 113, 10759–10768. doi:10.1073/pnas.1605516113

Nottingham, A.T., Fierer, N., Turner, B.L., Whitaker, J., Ostle, N.J., McNamara, N.P., Bardgett, R.D., Leff, J.W., Salinas, N., Silman, M.R., 2018. Microbes follow Humboldt: temperature drives plant and soil microbial diversity patterns from the Amazon to the Andes. Ecology 99, 2455–2466.

Oksanen, J., Blanchet, F.G., Friendly, M., Kindt, R., Legendre, P., McGlinn, D., Minchin, P.R., O’hara, R.B., Simpson, G.L., Solymos, P., 2016. vegan: Community Ecology Package. R package version 2.4-3. Vienna: R Foundation for Statistical Computing.[Google Scholar].

Parada, A.E., Needham, D.M., Fuhrman, J.A., 2016. Every base matters: assessing small subunit rRNA primers for marine microbiomes with mock communities, time series and global field samples. Environmental Microbiology 18, 1403–1414. doi:10.1111/1462-2920.13023

Pascault, N., Ranjard, L., Kaisermann, A., Bachar, D., Christen, R., Terrat, S., Mathieu, O., Lévêque, J., Mougel, C., Henault, C., 2013. Stimulation of different functional groups of bacteria by various plant residues as a driver of soil priming effect. Ecosystems 16, 810– 822.

Paula, F.S., Rodrigues, J.L.M., Zhou, J., Wu, L., Mueller, R.C., Mirza, B.S., Bohannan, B.J.M., Nüsslein, K., Deng, Y., Tiedje, J.M., 2014. Land use change alters functional gene diversity, composition and abundance in Amazon forest soil microbial communities. Molecular Ecology 23, 2988–2999.

Pedrinho, A., Mendes, L.W., Merloti, L.F., da Fonseca, M. de C., Cannavan, F. de S., Tsai, S.M., 2019. Forest-to-pasture conversion and recovery based on assessment of microbial communities in Eastern Amazon rainforest. FEMS Microbiology Ecology 95. doi:10.1093/femsec/fiy236

Peres-Neto, P.R., Jackson, D.A., 2001. How well do multivariate data sets match? The advantages of a Procrustean superimposition approach over the Mantel test. Oecologia 129, 169–178. doi:10.1007/s004420100720

Petersen, I.A.B., Meyer, K.M., Bohannan, B.J.M., 2019. Meta-analysis reveals consistent bacterial responses to land use change across the tropics. Frontiers in Ecology and Evolution 7, 391.

Ponge, J.-F., 2015. The soil as an ecosystem. Biology and Fertility of Soils 51, 645–648.

Poorter, L., van der Sande, M.T., Thompson, J., Arets, E.J.M.M., Alarcón, A., Álvarez-Sánchez, J., Ascarrunz, N., Balvanera, P., Barajas-Guzmán, G., Boit, A., Bongers, F., Carvalho, F.A., Casanoves, F., Cornejo-Tenorio, G., Costa, F.R.C., de Castilho, C. V, Duivenvoorden, J.F., Dutrieux, L.P., Enquist, B.J., Fernández-Méndez, F., Finegan, B., Gormley, L.H.L., Healey, J.R., Hoosbeek, M.R., Ibarra-Manríquez, G., Junqueira, A.B., Levis, C., Licona, J.C., Lisboa, L.S., Magnusson, W.E., Martínez-Ramos, M., Martínez-Yrizar, A., Martorano, L.G., Maskell, L.C., Mazzei, L., Meave, J.A., Mora, F., Muñoz, R., Nytch, C., Pansonato, M.P., Parr, T.W., Paz, H., Pérez-García, E.A., Rentería, L.Y., Rodríguez-Velazquez, J., Rozendaal, D.M.A., Ruschel, A.R., Sakschewski, B., Salgado-Negret, B., Schietti, J., Simões, M., Sinclair, F.L., Souza, P.F., Souza, F.C., Stropp, J., ter Steege, H., Swenson, N.G., Thonicke, K., Toledo, M., Uriarte, M., van der Hout, P., Walker, P., Zamora, N., Peña-Claros, M., 2015. Diversity enhances carbon storage in tropical forests. Global Ecology and Biogeography 24, 1314–1328. doi:https://doi.org/10.1111/geb.12364

Prober, S.M., Leff, J.W., Bates, S.T., Borer, E.T., Firn, J., Harpole, W.S., Lind, E.M., Seabloom, E.W., Adler, P.B., Bakker, J.D., 2015. Plant diversity predicts beta but not alpha diversity of soil microbes across grasslands worldwide. Ecology Letters 18, 85–95.

Purahong, W., Wubet, T., Lentendu, G., Schloter, M., Pecyna, M.J., Kapturska, D., Hofrichter, M., Krüger, D., Buscot, F., 2016. Life in leaf litter: novel insights into community dynamics of bacteria and fungi during litter decomposition. Molecular Ecology 25, 4059–4074.

Quast, C., Pruesse, E., Yilmaz, P., Gerken, J., Schweer, T., Yarza, P., Peplies, J., Glöckner, F.O., 2012. The SILVA ribosomal RNA gene database project: improved data processing and web-based tools. Nucleic Acids Research 41, D590–D596.

Rillig, M.C., Ryo, M., Lehmann, A., Aguilar-Trigueros, C.A., Buchert, S., Wulf, A., Iwasaki, A., Roy, J., Yang, G., 2019. The role of multiple global change factors in driving soil functions and microbial biodiversity. Science 366, 886–890.

Ritter, C.D., Zizka, A., Roger, F., Tuomisto, H., Barnes, C., Nilsson, R.H., Antonelli, A., 2018. High-throughput metabarcoding reveals the effect of physicochemical soil properties on soil and litter biodiversity and community turnover across Amazonia. PeerJ 6, e5661. doi:10.7717/peerj.5661

Rodrigues, J.L.M., Pellizari, V.H., Mueller, R., Baek, K., Jesus, E. da C., Paula, F.S., Mirza, B., Hamaoui, G.S.J., Tsai, S.M., Feigl, B., Tiedje, J.M., Bohannan, B.J.M., Nüsslein, K., 2013. Conversion of the Amazon rainforest to agriculture results in biotic homogenization of soil bacterial communities. Proceedings of the National Academy of Sciences of the United States of America 110, 988–993. doi:10.1073/pnas.1220608110

Santos Lemos, R.D. dos, R.C. de Santos H.G. dos, Ker Anjos, J.C., L.H.C. dos, Shimizu, S.H., 2005. Manual de descrição e coleta de solo no campo.

Sayer, E.J., Rodtassana, C., Sheldrake, M., Bréchet, L.M., Ashford, O.S., Lopez-Sangil, L., Kerdraon-Byrne, D., Castro, B., Turner, B.L., Wright, S.J., 2020. Revisiting nutrient cycling by litterfall—Insights from 15 years of litter manipulation in old-growth lowland tropical forest, in: Advances in Ecological Research. Elsevier, pp. 173–223.

Sayer, E.J., Tanner, E.V.J., 2010. Experimental investigation of the importance of litterfall in lowland semi-evergreen tropical forest nutrient cycling. Journal of Ecology 98, 1052–1062. doi:10.1111/j.1365-2745.2010.01680.x

Segata, N., Izard, J., Waldron, L., Gevers, D., Miropolsky, L., Garrett, W.S., Huttenhower, C., 2011. Metagenomic biomarker discovery and explanation. Genome Biology 12, 1–18.

Souza, J.L.L. de S., Fontes, M.P.F., Gilkes, R., Costa, L.M. da, Oliveira, T.S. de, 2018. Geochemical Signature of Amazon Tropical Rainforest Soils. Revista Brasileira de Ciência do Solo 42.

Suleiman, A.K.A., Manoeli, L., Boldo, J.T., Pereira, M.G., Roesch, L.F.W., 2013. Shifts in soil bacterial community after eight years of land-use change. Systematic and Applied Microbiology 36, 137–144. doi:https://doi.org/10.1016/j.syapm.2012.10.007

Tate, K.R., 2015. Soil methane oxidation and land-use change–from process to mitigation. Soil Biology and Biochemistry 80, 260–272.

Team, R.C., 2018. R: A language and environment for statistical computing. R Foundation for Statistical Computing, Vienna, Austria. Version 3.6. 1.

Teixeira, P.C., Donagemma, G.K., Fontana, A., Teixeira, W.G., 2017. Manual de métodos de análise de solo. Rio de Janeiro, Embrapa. 573p.

Tláskal, V., Voříšková, J., Baldrian, P., 2016. Bacterial succession on decomposing leaf litter exhibits a specific occurrence pattern of cellulolytic taxa and potential decomposers of fungal mycelia. FEMS Microbiology Ecology 92, fiw177.

Tripathi, B.M., Stegen, J.C., Kim, M., Dong, K., Adams, J.M., Lee, Y.K., 2018. Soil pH mediates the balance between stochastic and deterministic assembly of bacteria. The ISME Journal 12, 1072–1083.

Ushio, M., Kitayama, K., Balser, T.C., 2010. Tree species-mediated spatial patchiness of the composition of microbial community and physicochemical properties in the topsoils of a tropical montane forest. Soil Biology and Biochemistry 42, 1588–1595. doi:https://doi.org/10.1016/j.soilbio.2010.05.035

Ventura, M., Canchaya, C., Tauch, A., Chandra, G., Fitzgerald, G.F., Chater, K.F., van Sinderen, D., 2007. Genomics of Actinobacteria: tracing the evolutionary history of an ancient phylum. Microbiology and Molecular Biology Reviews 71, 495–548.

Wright, E.S., Yilmaz, L.S., Noguera, D.R., 2012. DECIPHER, a search-based approach to chimera identification for 16S rRNA sequences. Applied and Environmental Microbiology 78, 717–725. doi:10.1128/AEM.06516-11

Xue, B., Xie, J., Huang, J., Chen, L., Gao, L., Ou, S., Wang, Y., Peng, X., 2016. Plant polyphenols alter a pathway of energy metabolism by inhibiting fecal Bacteroidetes and Firmicutes in vitro. Food & Function 7, 1501–1507.

Yamada, T., Sekiguchi, Y., Imachi, H., Kamagata, Y., Ohashi, A., Harada, H., 2005. Diversity, localization, and physiological properties of filamentous microbes belonging to Chloroflexi subphylum I in mesophilic and thermophilic methanogenic sludge granules. Applied and Environmental Microbiology 71, 7493–7503.

Yuan, Z.Y., Chen, H.Y.H., 2010. Fine root biomass, production, turnover rates, and nutrient contents in boreal forest ecosystems in relation to species, climate, fertility, and stand age: literature review and meta-analyses. Critical Reviews in Plant Sciences 29, 204–221.

Zechmeister-Boltenstern, S., Keiblinger, K.M., Mooshammer, M., Peñuelas, J., Richter, A., Sardans, J., Wanek, W., 2015. The application of ecological stoichiometry to plant– microbial–soil organic matter transformations. Ecological Monographs 85, 133–155.

