## Supplemental information for "Land-use and forest floor explain prokaryotic metacommunity structuring and spatial turnover in Amazonian forest-to-pasture conversion areas"

**This PDF file includes:**

Supplementary text

Figure S1 to S7

Table S1 to S6

Supplemental information Text

**Soil variable selection**

Principal component analysis (PCA) was used to compare samples originating from the studied soil fertility gradient across the sites. The data was standardized using *(xi – mean(x))/sd(x)*, where *mean(x)* is the mean of x values, and *sd(x)* is the standard deviation (SD). The total contribution of a given variable on principal component axes was estimated. For example, the observed contributions of a variable on two principal components, say PC1 and PC2, was calculated with the formula: *[(C1 * Eig1) + (C2 * Eig2)] / (Eig1 + Eig2)*, where C1 and C2 are the contributions of the variable to PC1 and PC2, and Eig1 and Eig2 the eigenvalues of PC1 and PC2, respectively. The expected average contribution of a variable to PC1 and PC2 is: *[(number of variables * Eig1) + (number of variables * Eig2)] / (Eig1 + Eig2)*. In this study, the expected value was 1/length(variables) = 1/20 = 5%. In our results, variables with a contribution larger than this cutoff were considered as important in contributing to associated components. The visualization of the PCA was created with the *PCA()* function of the 'FactoMiner' R package v.2.3 (Husson et al., 2017).

**Selected soil variables and correspondence in the structuring of soil microbial communities**

Variables were selected based on their explained variance of the PCA. Next, a Constrained Analysis of Principal coordinates (CAP) was performed with these variables and a matrix of Bray-Curtis distances (*capscale (formula = distance ~ silt + pH + %BS +* $Al^{+3}$ *saturation +* $Ca^{+2}$*+*$Mg^{+2}$*, data = data)*) using the *ordinate()* function of 'phyloseq' v.1.30.0 (McMurdie and Holmes, 2013). All selected soil variables revealed a significant relationship with the structure of the prokaryotic metacommunity (i.e., assemblage of communities) and estimated significance of correlation between soil variables and observed differences between samples.

**Diversity partitioning analysis**

Diversity profiles (i.e., alpha (α), beta (β), and gamma (γ) diversities) were calculated according to Hill numbers (for more details see Chao et al. (2014)). The ASV richness, the exponential of Shannon's entropy, and the Simpson index were calculated by giving the values 0, 1, and 2 for the Hill parameter q, respectively. When q = 0, the Hill index is sensitive to ASV abundance. All individuals are equally weighted when q = 1, and the index is insensitive to the dominant species when q = 2. This approach was used to evaluate the contribution of each compartment and the overall forest floor (i.e., litter, root layer, and bulk soil) for each diversity scale. The package 'entropart' v.1.6.1 (Marcon and Hérault, 2015) was used. Specifically, the *DivProfile()* function was used to calculate alpha, beta and gamma diversities. The *MergeMC()* function was used to aggregate the data present in each of the compartments into a single object, which made it possible to extract the diversity measures for the forest floor.

Finally, to test our hypothesis that the alpha, beta, and gamma diversities of the forest compartments combined are higher than in that of the pasture soil, we used the Kruskal-Wallis test to detect statistical differences. Prokaryotic diversity was compared between the forest and pasture soils as well as between the forest floor and pasture soil. The comparison was based on the overall effective numbers (i.e., Hill's q 0, 1 and 2). All p-values were corrected for false discovery rate (Benjamini-Hochberg FDR correction).

**References**

Chao, A., Chiu, C.-H., Jost, L., 2014. Unifying species diversity, phylogenetic diversity, functional diversity, and related similarity and differentiation measures through Hill numbers. Annual Review of Ecology, Evolution, and Systematics 45, 297–324.

Husson, F., Lê, S., Pagès, J., 2017. Exploratory multivariate analysis by example using R. CRC press.

Marcon, E., Hérault, B., 2015. entropart: An R package to measure and partition diversity. Journal of Statistical Software 67, 1–26.

McMurdie, P.J., Holmes, S., 2013. phyloseq: an R package for reproducible interactive analysis and graphics of microbiome census data. PloS One 8, e61217. doi:10.1371/journal.pone.0061217

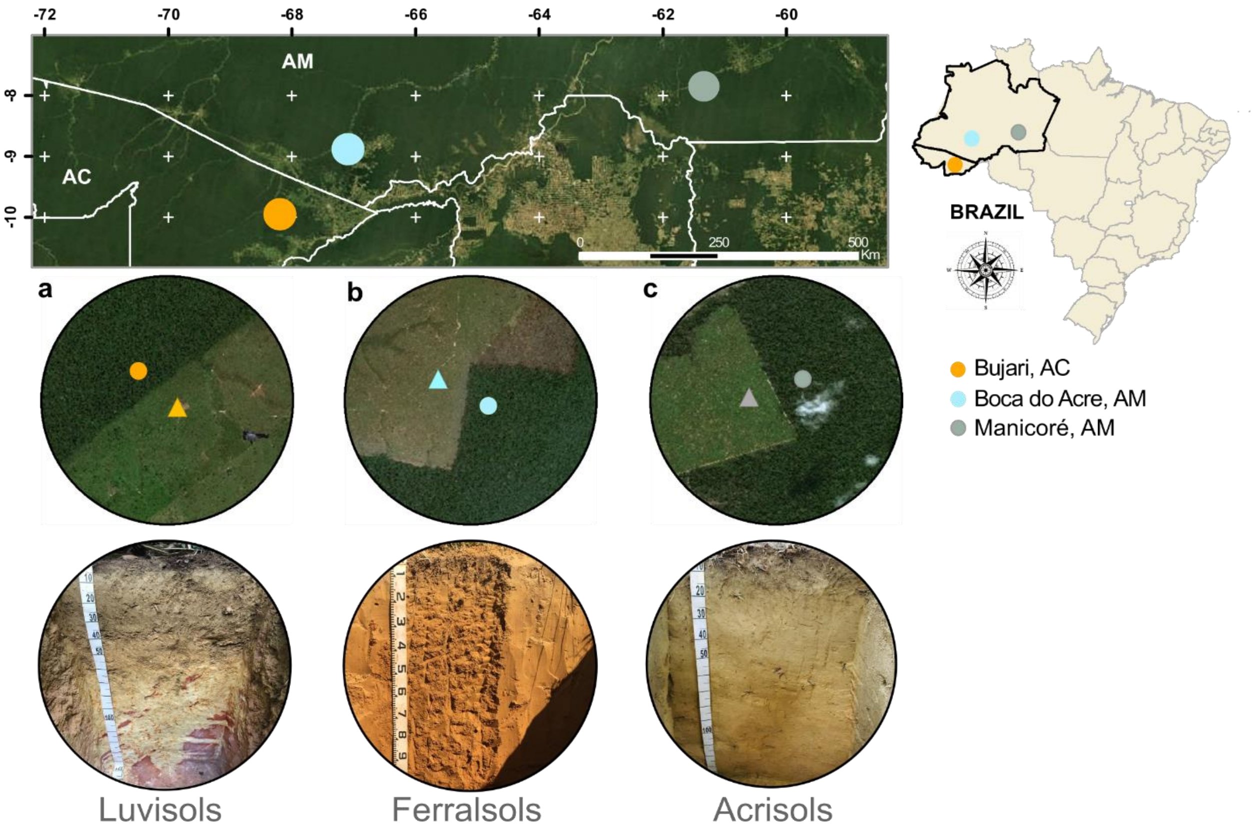


**Figure S1. Study area and sampling locations.** Map of the study area in the Brazilian Western Amazonia, with forest-to-pasture conversion areas and their predominant soil orders shown below. **(a)** Bujari (BUJ), state of Acre (AC); **(b)** Boca do Acre (BAC), state of Amazonas (AM); and **(c)** Manicoré (MAN) also in the state of Amazonas (AM). Satellite images provided by ESRI Service Layer.

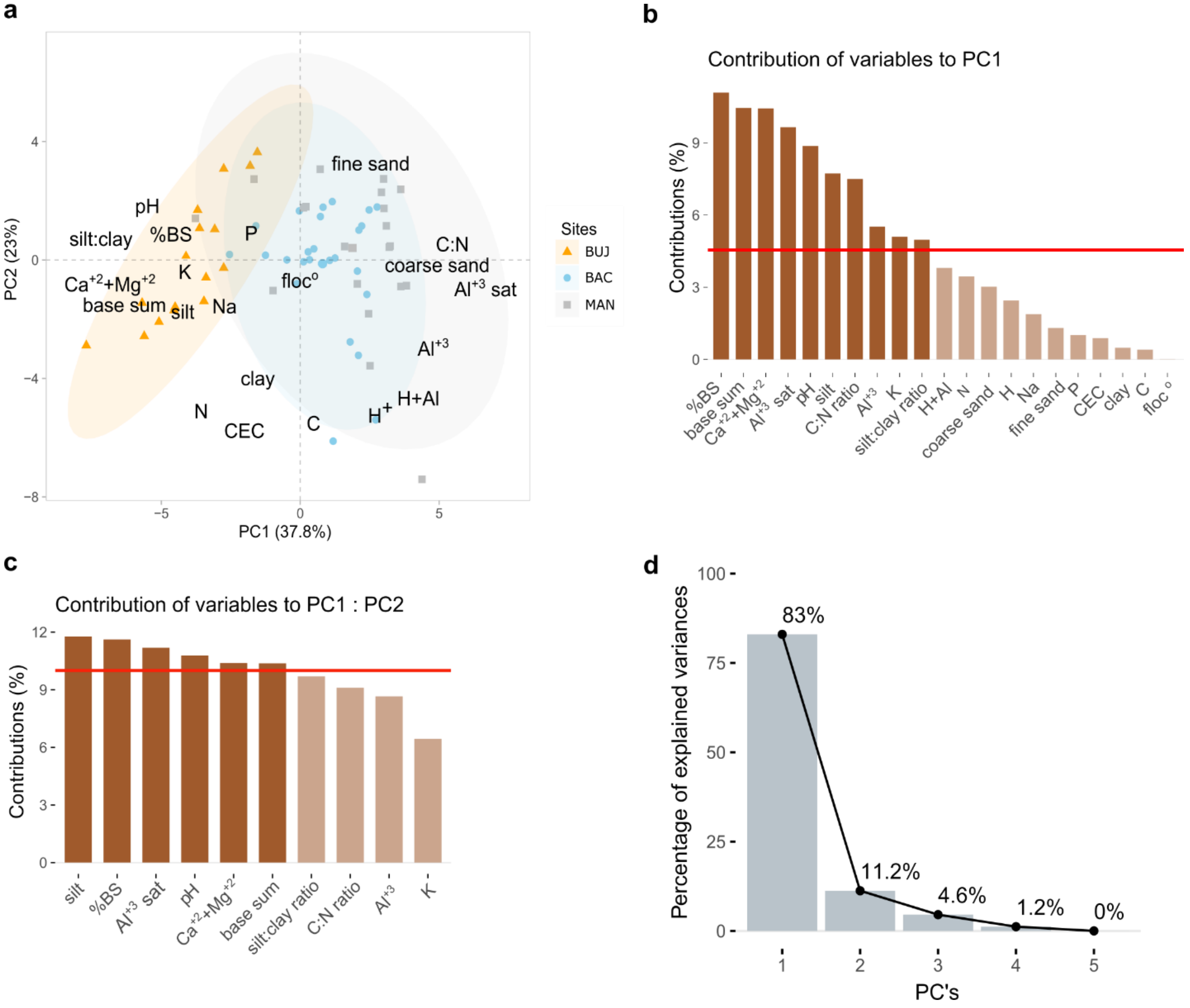


**Figure S2. Soil variable selection.** A principal component analysis (PCA) of soil chemical and physical variables was calculated **(a)**, and the variables that contributed to PC1 above a given threshold (red line) **(b)** were selected and used for a new PCA. They went through a further selection, now based on their loadings on both PC1 and PC2 **(c)**. The remaining variables explained a large amount of the explained variances (%), showing the first axis as the most important to detect the gradient of fertility among the study sites **(d)**.

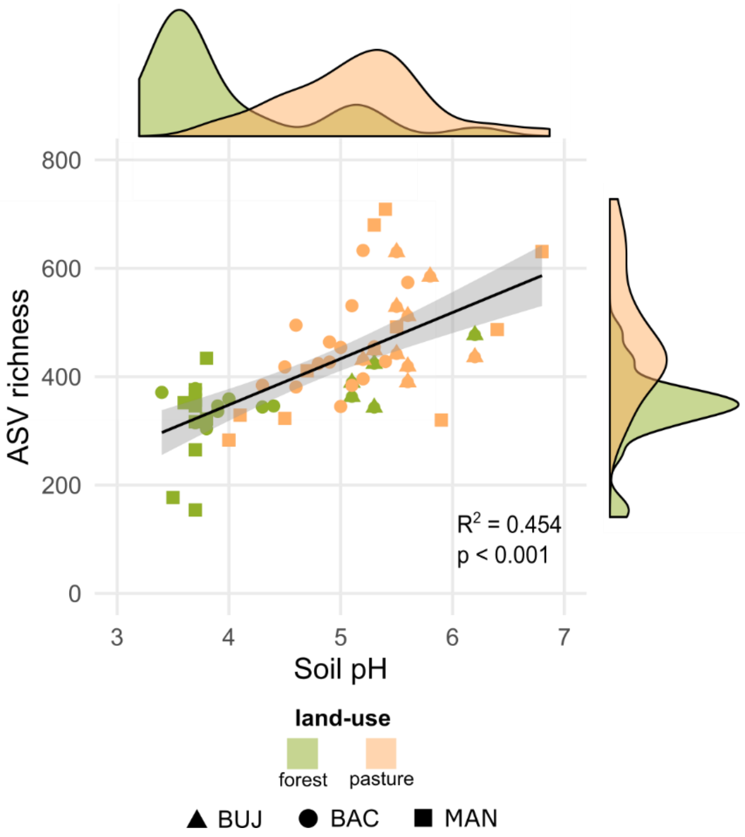


**Figure S3. Soil pH effects on the ASV richness in forest-to-pasture conversion sites of the Western Brazilian Amazonia.** Correlation between ASV richness and soil pH of each evaluated site in both land uses. The fitted values for each model are represented by the black line and their standard errors are indicated by the shaded area.

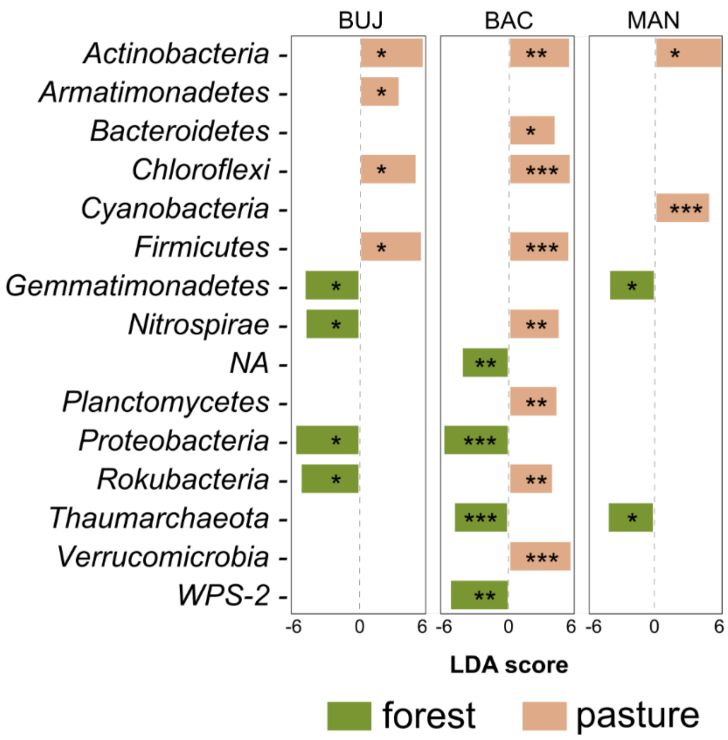


**Figure S4. Soil prokaryotes with significantly different abundances between land uses.** LefSe multivariate analysis (false discovery rate (FDR) adjusted p-value < 0.05, LDA > 2.0) between land uses in Bujari (BUJ), Boca do Acre (BAC) and Manicoré (MAN).

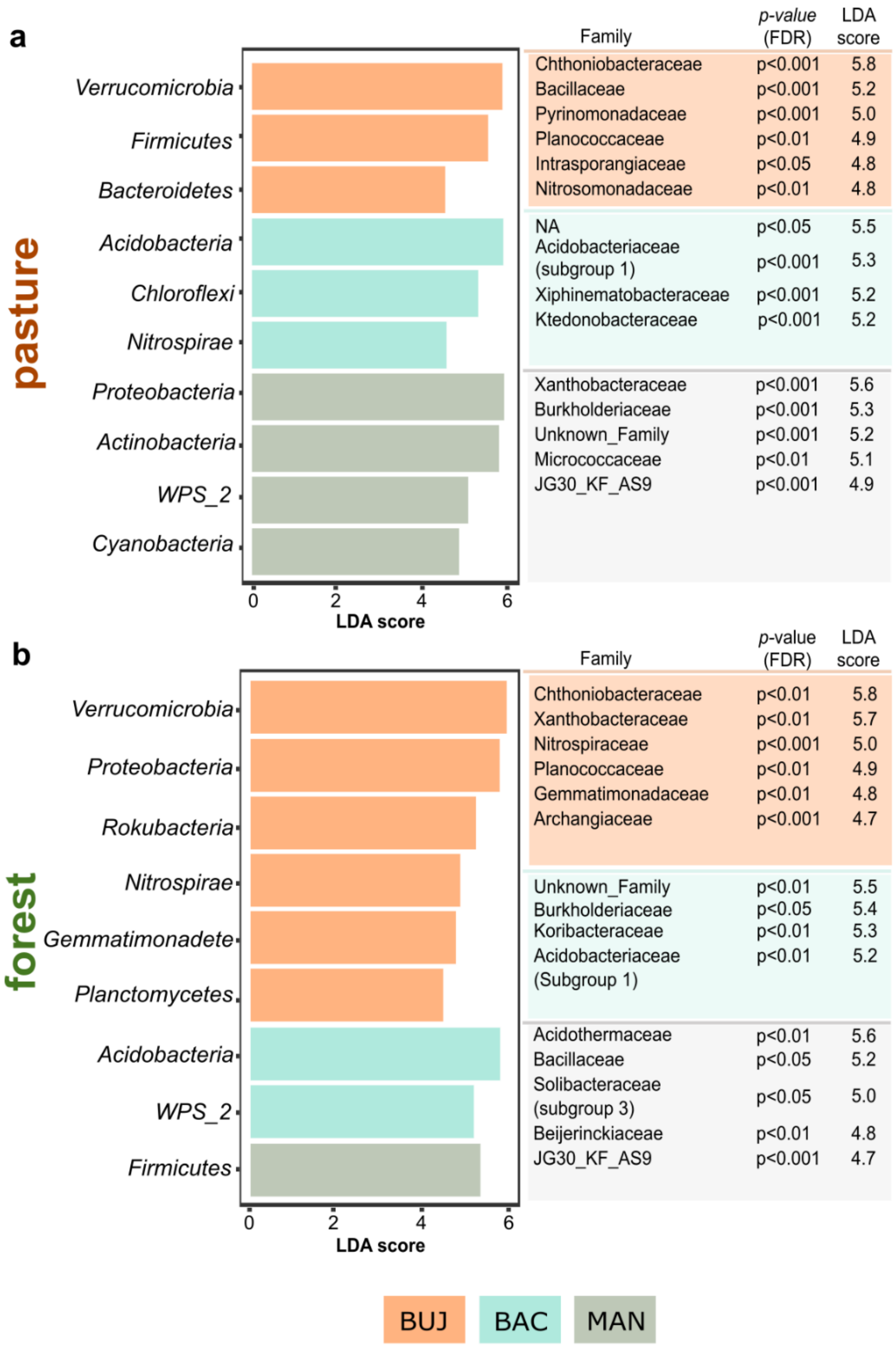


**Figure S5. Soil prokaryotes with significantly different abundances within land uses across study sites.** LefSe multivariate analysis (false discovery rate (FDR) adjusted p-value < 0.05, LDA > 2.0). Comparisons of the prokaryotic communities (phylum and family levels) between sites in the **(a)** pasture and **(b)** forest.

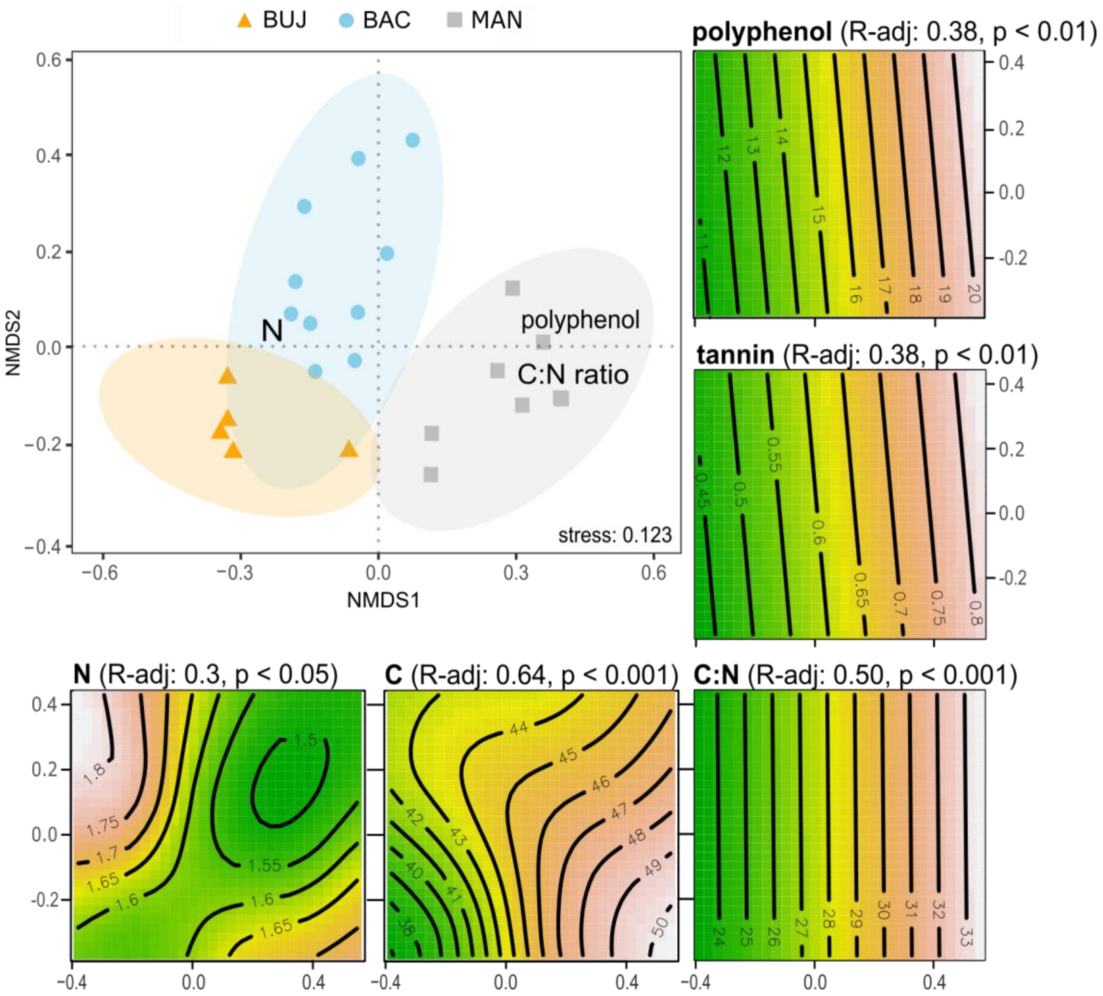


**Figure S6. Prokaryotic community structure in the litter varies with litter chemistry in Brazilian Western Amazonian forests.** Nonmetric multidimensional scaling **(**NMDS) based on a Bray-Curtis dissimilarity matrix showing variation in community structure in function of the litter chemical variables and tested by generalized additive models (GAM).

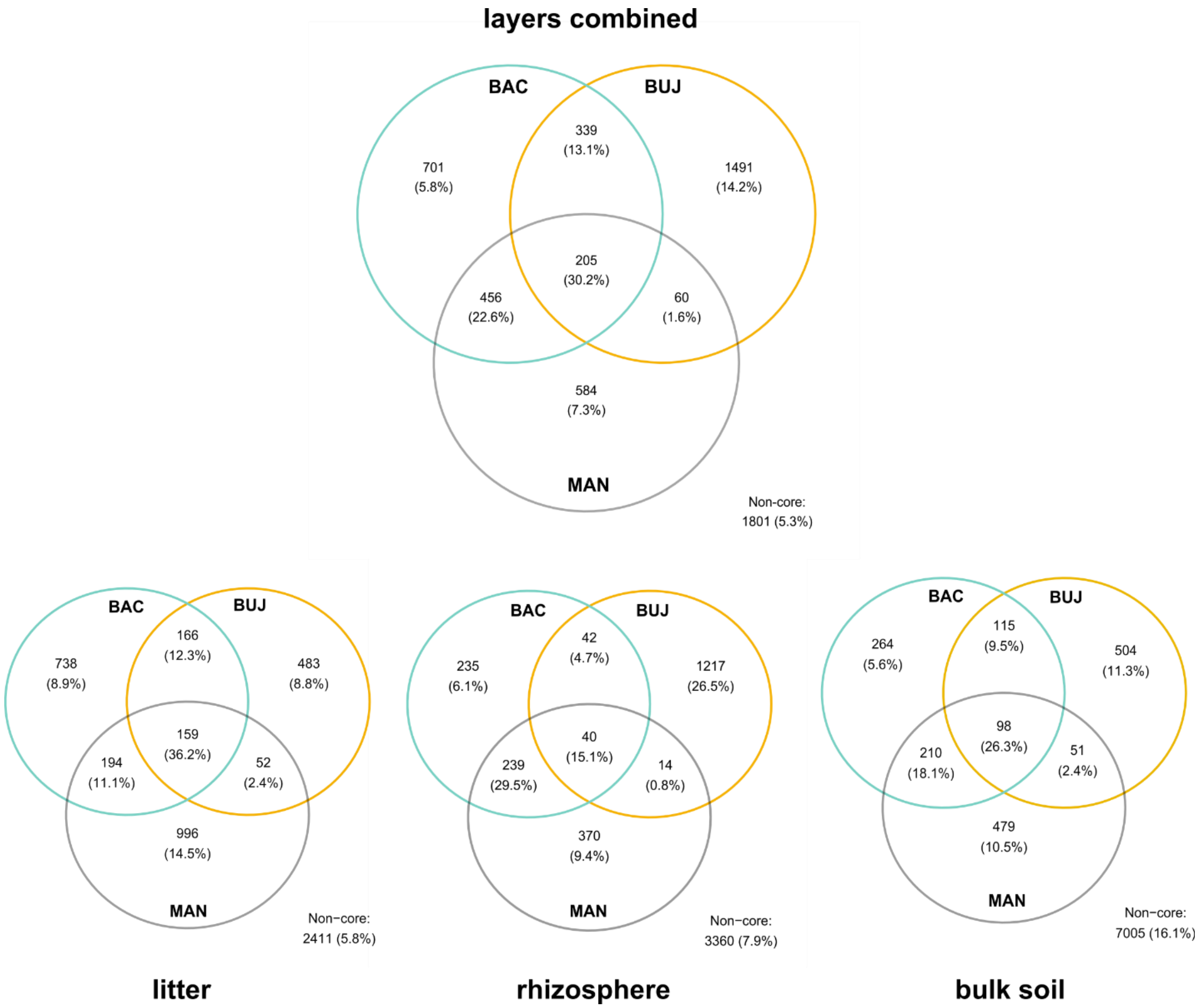


**Figure S7. ASVs shared between regions and forest floor compartments.** Venn diagram for each compartment of the forest floor, showing the most frequent (observed in at least 10% of the samples) and abundant (relative abundance in each sample greater than 0.01%) ASVs.

**Table S1.** Soil and landscape features, pedological descriptions, and geographic coordinates of the regions and land-use systems selected in the Brazilian Western Amazonia.

**See supplementary spreadsheet (.xlsx)**

**Table S2.** Kruskal-Wallis chi-squared test (Bonferroni corrected p-values) comparing selected soil variables between land uses (forest x pasture), forests (between forests by sites) and pastures (between pastures by sites).

| Source of variation | Comparison | silt | BS% | $Al^{+3}$ saturation | pH | Ca+Mg | base sum |
|---|---|---|---|---|---|---|---|
| | | $\chi^2 = 51.62$ | $\chi^2 = 46.86$ | $\chi^2 = 45.34$ | $\chi^2 = 43.37$ | $\chi^2 = 49.96$ | $\chi^2 = 49.59$ |
| Land-use within | BUJ | n.s. | n.s. | n.s. | n.s. | n.s. | n.s. |
| | BAC | n.s. | 0.001 | 0.001 | 0.001 | 0.001 | 0.001 |
| | MAN | n.s. | 0.001 | 0.001 | 0.001 | 0.002 | 0.001 |
| Forest between | BUJ x BAC | 0.007 | 0.002 | 0.003 | 0.002 | 0.003 | 0.002 |
| | BAC x MAN | 0.001 | 0.006 | n.s. | 0.001 | n.s. | 0.001 |
| | BUJ x MAN | 0.002 | 0.002 | 0.001 | 0.001 | 0.002 | 0.001 |
| Pasture between | BUJ x BAC | 0.001 | 0.001 | 0.001 | 0.001 | 0.001 | 0.001 |
| | BAC x MAN | 0.001 | n.s. | n.s. | n.s. | n.s. | n.s. |
| | BUJ x MAN | 0.001 | 0.007 | n.s. | 0.001 | n.s. | 0.001 |

**Table S3.** PERMANOVA testing the effects of land-use and sites on the bulk soil prokaryotic metacommunity; Land-use (forest x pasture), forest (between forests by sites) and pasture (between pastures by sites).

| Source of variation | Comparisons | F | p-value |
|---|---|---|---|
| | BUJ | 4.86 | 0.001 |
| Land-use within | BAC | 9.67 | 0.001 |
| | MAN | 3.60 | 0.002 |
| | BUJ xBAC | 9.93 | 0.001 |
| Forest between | BAC xMAN | 8.11 | 0.001 |
| | BUJ xMAN | 3.72 | 0.001 |
| | BUJ xBAC | 7.81 | 0.001 |
| Pasture between | BAC xMAN | 7.52 | 0.001 |
| | BUJ xMAN | 6.06 | 0.001 |
| (interaction) | sites xland-use | 3.97 | 0.001 |

**Table S4.** Generalized Additive Model outputs testing the correlation of each of the selected soil variables on the soil prokaryotic metacommunity structuring.

| | | Deviance explained (%) | F | p-value |
|---|---|---|---|---|
| Soil chemical variables | pH | 91.6 | 61.92 | 0.001 |
| | % BS | 93.9 | 88.28 | 0.001 |
| | $Al^{+3}$ saturation | 95.2 | 112.7 | 0.001 |
| | Ca + Mg | 80.2 | 22.62 | 0.001 |
| Soil physical variable | silt | 76.9 | 18751 | 0.001 |

**Table S5.** PERMANOVA testing the effect of sites on the litter and root layer prokaryotic metacommunity.

| Source of variation | Comparisons | F | p-value |
|---|---|---|---|
| Litter between | BUJ xBAC | 2.69 | 0.002 |
| | BAC xMAN | 4.82 | 0.001 |
| | BUJ xMAN | 4.10 | 0.001 |
| Root layer between | BUJ xBAC | 15.94 | 0.001 |
| | BAC xMAN | 15.04 | 0.001 |
| | BUJ xMAN | 9.75 | 0.001 |

**Table S6.** Kruskal-Wallis chi-squared test for scales of diversity between the forest floor (litter, root layer, and bulk soil) and pasture bulk soil.

| | α - diversity | | β - diversity | | γ - diversity | |
|---|---|---|---|---|---|---|
| | $\chi^2$ | p-value | $\chi^2$ | p-value | $\chi^2$ | p-value |
| BUJ | 0.109 | 0.743 | 16.916 | < **0.001** | 6.648 | **0.009** |
| BAC | 6.607 | **0.014** | 9.967 | < **0.01** | 0.411 | 0.521 |
| MAN | 2.632 | 0.104 | 12.405 | < **0.001** | 0.172 | 0.678 |
